# Humanization of antibodies using a machine learning approach on large-scale repertoire data

**DOI:** 10.1101/2021.01.08.425894

**Authors:** Mark Chin, Claire Marks, Charlotte M. Deane

## Abstract

Monoclonal antibody therapeutics are often produced from non-human sources (typically murine), and can therefore generate immunogenic responses in humans. Humanization procedures aim to produce antibody therapeutics that do not elicit an immune response and are safe for human use, without impacting efficacy. Humanization is normally carried out in a largely trial-and-error experimental process. We have built machine learning classifiers that can discriminate between human and non-human antibody variable domain sequences using the large amount of repertoire data now available. Our classifiers consistently outperform existing best-in-class models, and our output scores exhibit a negative relationship with the experimental immunogenicity of existing antibody therapeutics. We used our classifiers to develop a novel, computational humanization tool, Hu-mAb, that suggests mutations to an input sequence to reduce its immunogenicity. For a set of existing therapeutics with known precursor sequences, the mutations suggested by Hu-mAb show significant overlap with those deduced experimentally. Hu-mAb is therefore an effective replacement for trial-and-error humanization experiments, producing similar results in a fraction of the time. Hu-mAb is freely available to use at opig.stats.ox.ac.uk/webapps/humab.

## Introduction

Since the first monoclonal antibody (mAb), muromonab, was approved by the US FDA in 1986, the antibody therapeutic market has grown exponentially, with six of the top ten selling drugs in 2018 being mAbs (1). Therapeutic mAbs and antibody-related products such as Fc-fusion proteins, antibody fragments, nanobodies, and antibody-drug conjugates are now the predominant class of biopharmaceuticals, representing half the total sales of all biopharmaceutical products (2). These therapeutics treat a range of pathologies including but not limited to cancer, multiple sclerosis, asthma and rheumatoid arthritis. To date (September 2020), 93 therapeutics mAbs have been approved by the US FDA and at least 400 others are in development (3).

Many therapeutic antibodies are derived from natural B-cell repertoires of mice, or mice with an engineered human germline repertoire (1). However, antibodies developed in animal models are often not tolerated by humans and can elicit an immune response – this property is known as immunogenicity. Immunogenic responses can negatively impact both safety and pharmacokinetic properties of the therapeutics and can result in the production of neutralizing antibodies that lead to loss of efficacy (1). This can pose a significant barrier to the development and approval of therapeutics (4). To combat the immunogenicity of mAbs, various techniques to engineer murine antibodies by substituting part of their sequence with human ones are used. These include chimerization (5) and humanization (6). The former involves the combining of a murine variable domain with human constant region domains, and the latter involves grafting the murine CDR sequences into a human scaffold. Early studies have suggested that more human-like sequences demonstrate lower levels of immunogenicity (7). Whilst multiple techniques have been developed to obtain fully human mAbs, humanized antibodies remain the predominant class of mAb making up 50% of therapeutics in development (3).

The aim of humanization is to reduce immunogenicity while preserving the efficacy of the therapeutic. Typically, human frameworks with high homology to the original sequence of interest are chosen as a scaffold (8). Some murine residues in framework regions, referred to as vernier zone residues, affect the conformation of CDR loops and may therefore be retained to preserve antibody affinity. Although computational methods are available (e.g. 9–14), the humanization process remains a bottleneck in mAb development, often based on trial-and-error, involving arbitrary back-mutations to restore efficacy or reduce immunogenicity (15).

An effective systematic humanization protocol requires a model that is able to identify the humanness of a sequence with little error. Higher humanness scores should also be linked with lower levels of immunogenicity. Traditional humanness scores are based on pairwise sequence identity methods between the sample and a set of reference (most often germline) human sequences. For example, the score can correspond to the sequence identity of the closest germline sequence or the average among a set of sequences (11). Recent models take account of both preferences of particular residues and pair correlations between amino acids (12). A Multivariate Gaussian model (MG) utilized a statistical inference approach (13) and could distinguish human from murine sequences accurately, but the score demonstrated only a weak negative correlation to experimental immunogenicity levels. More recently, a deep learning approach utilizing a bidirectional long short-term memory (LSTM) model demonstrated best-in-class performance in discriminating between human and murine sequences (14).

The recent growth of publicly available antibody sequences has created many opportunities for large-scale data mining. OAS (16) is a database of Ig-seq outputs from *∼*80 studies with nearly 2 billion redundant antibody sequences across diverse immune states and organisms (although primarily human and mouse). OAS is ideal for data mining due to its size, consistent IMGT numbering, and because the sequences represent natural mature antibodies produced *in vivo*.

Utilizing random forest models, we have constructed classifiers that accurately distinguish between each human V gene and non-human variable domain sequences. The ‘human-ness’ scores produced by these classifiers exhibited a negative relationship with observed immunogenicity levels. We used these models to build Hu-mAb – a computational tool that can systematically humanize VL and VH sequences of interest by suggesting mutations that increase humanness. Hu-mAb humanizes the sequence in an optimal manner, minimizing the number of mutations made to the sequence to limit the impact on efficacy. The mutations made by our humanizer were found to be very similar to those made in experimental therapeutic humanization studies that produced sequences with low immunogenicity. Hu-mAb offers a powerful alternative to time-consuming, trial-and-error based approaches to reducing immunogenicity. Hu-mAb is freely available at opig.stats.ox.ac.uk/webapps/humab.

## Results

### Classification performance of our Random Forest (RF) models on OAS sequences

RF models were generated by training on the OAS IgG dataset (see Methods). Each model was created as a binary classifier – trained on human antibody sequences (either VH, VL Kappa or VL Lambda) of a specific V gene type as the positive class and all non-human sequences of the respective chain type as the negative class. Different classifiers were constructed for each V gene as PCA demonstrated trivial clustering of sequences by their respective V gene type (see SI, section 3A). The performance of the RF models was assessed by determining its ability to correctly distinguish human sequences of a specific V gene type from those originating from other species. We used the validation set to determine the classification thresholds as the value that maximizes the Youden’s J statistic (YJS; see Methods). Performance on the test set was then calculated using the chosen threshold for each model. Extremely high performance was observed across all models, achieving AUCs (area under the receiver operating characteristic curve) close to 1 or 1 (see SI, section 3B). Similar YJS values were also seen in both validation and test sets with all models scoring *≥*0.999. All the VH models perfectly discriminated between human and negative sequence in both validation and test sets. Performance on the light chain is slightly worse - this may be due to the greater amount of negative training data available for the VH models (>12 million sequences) compared to that of kappa (950,000) and lambda (650,000) models.

### Comparison of RF models to previous LSTM models

Recent work has used the LSTM model for predicting humanness (14). We generated LSTM models with our dataset of sequences (see Methods) and performance was compared to our RF models. Across all 22 models (each chain and each V gene type), the RF model outperformed the respective LSTM model on both AUC and YJS scores (see SI, section 3C). None of the LSTM models were capable of completely discriminating between human and negative sequences. We suspect our RF models produce better results because they are trained using both positive and negative (non-human) data, whereas LSTM models were only trained on positive human sequences.

### Classification of therapeutics

A set of 481 antibody therapeutics (Phase I to approved) were obtained from Thera-SAbDab (3) (see Methods). Each VH and VL sequence was scored by the respective set of RF classifiers (VH, kappa, or lambda) and was classified as human if a single model scores it as human (above the YJS threshold). In the case of VL sequences, we built and used an additional RF model to first discriminate whether the sequence type is kappa or lambda (see Methods). Figure 1 shows the proportion of therapeutics classified as human (split by origin) by their chain type (VH or VL) and combined (requires both VH and VL to be classified as human). All 176 human sequences were classified as human, and all 14 mouse sequences as non-human. Overall our RF models classify more therapeutics as human, as the human content of the antibody sequences increases. This trend is also observed using the LSTM method, but not as clearly - for example, more of the chi/humanized set are classified as human than the humanized set, and some human therapeutics are classified as non-human (SI section 3D).

**Fig 1.**
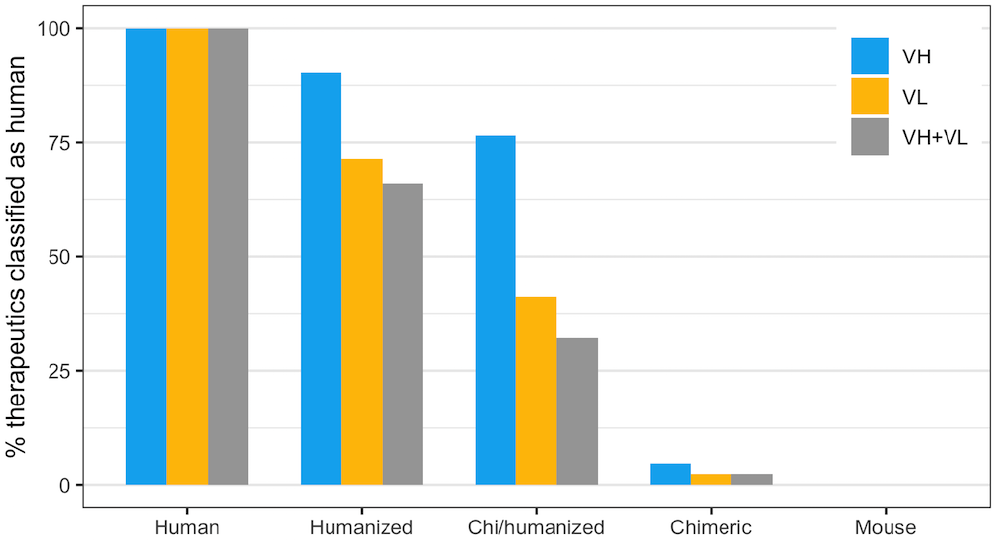
Percentage of antibody therapeutics classified as human by our RF models, split by their origin. Chi/humanized are sequences which are part humanized and part chimeric. Therapeutics were classified based on their VH and VL sequences separately, as well as combined (to be classified as human, both VH and VL scores had to be above the respective YJS threshold). As the humanness of the therapeutics decreases (left to right), the proportion classified as human also decreases.

It might be expected that all chimeric sequences have a completely non-human variable domain as only the constant domains are replaced with human sequence. However, two VH sequences and one VL sequence (out of 43) were labelled as human by our classifiers. This is likely to be because these sequences were of *Macaca irus* origin – a species that was not present in the training dataset from the OAS. Two-thirds of the humanized therapeutics had both VH and VL classified as human. Humanized sequences often have arbitrary back mutations on the FR regions to improve efficacy which might explain why not all humanized sequences are classified as human. Moreover, the INN definition was changed in 2014 such that sequences with a chimeric origin could be given an INN that implied a humanized sequence (17). VL sequences had a lower proportion classified as human compared to VH sequences. This could be potentially attributed to the lower number of mutations made in VL sequences (on average 75% of the number of mutations made on VH sequences - see Table 1).

**Table 1.** Comparison between experimental humanization, and our computational tool Hu-mAb. The mutation ratio is the average number of mutations Hu-mAb suggested relative to the number of mutations made experimentally. The overlap ratio is the number of mutations that were both suggested by Hu-mAb and made experimentally, relative to the number of mutations suggested by the Hu-mAb. For the ‘unadjusted’ overlap ratio, only mutations to identical amino acid types were considered; the ‘adjusted’ version considers mutations to similar amino acid types to be a match (see SI section 2C).

| Therapeutic | Gene | VH |  |  |  |  | Gene | VL |  |  |  |  |
| --- | --- | --- | --- | --- | --- | --- | --- | --- | --- | --- | --- | --- |
|  |  | Unadjusted Overlap Ratio | Adjusted Overlap Ratio | # Hu-mAb Mutations | # Experimental Mutations | Mutation Ratio |  | Unadjusted Overlap Ratio | Adjusted Overlap Ratio | # Hu-mAb Mutations | # Experimental Mutations | Mutation Ratio |
| AntiCD28 | V3 | 63% | 79% | 19 | 33 | 58% | KV4 | 64% | 73% | 11 | 19 | 58% |
| Campath | V4 | 75% | 88% | 16 | 39 | 41% | KV1 | 67% | 67% | 3 | 14 | 21% |
| Bevacizumab | V3 | 50% | 57% | 14 | 25 | 56% | KV1 | 89% | 100% | 9 | 16 | 56% |
| Herceptin | V3 | 59% | 78% | 27 | 32 | 84% | KV1 | 88% | 88% | 8 | 22 | 36% |
| Omalizumab | V3 | 62% | 76% | 21 | 34 | 62% | KV1 | 89% | 95% | 19 | 25 | 76% |
| Eculizumab | V1 | 73% | 73% | 15 | 23 | 65% | KV1 | 83% | 83% | 12 | 20 | 60% |
| Tocilizumab | V4 | 64% | 86% | 14 | 23 | 61% | KV1 | 78% | 89% | 9 | 19 | 47% |
| Pembrolizumab | V1 | 73% | 73% | 11 | 23 | 48% | KV3 | 75% | 75% | 12 | 20 | 60% |
| Pertuzumab | V3 | 68% | 79% | 19 | 32 | 59% | KV1 | 80% | 90% | 10 | 20 | 50% |
| Ixekizumab | V1 | 75% | 75% | 12 | 29 | 41% | KV2 | 78% | 100% | 9 | 12 | 75% |
| Palivizumab | V2 | 75% | 83% | 12 | 18 | 67% | KV1 | 77% | 92% | 13 | 26 | 50% |
| Certolizumab | V3 | 61% | 78% | 18 | 31 | 58% | KV1 | 80% | 90% | 10 | 20 | 50% |
| Idarucizumab | V4 | 80% | 80% | 15 | 24 | 63% | KV2 | 67% | 67% | 6 | 8 | 75% |
| Reslizumab | V3 | 50% | 80% | 10 | 21 | 48% | KV1 | 83% | 100% | 6 | 20 | 30% |
| Solanezumab | V3 | 50% | 70% | 10 | 16 | 63% | KV2 | 88% | 100% | 8 | 10 | 80% |
| Lorvotuzumab | V3 | 90% | 90% | 10 | 13 | 77% | KV2 | 82% | 82% | 11 | 13 | 85% |
| Pinatuzumab | V3 | 61% | 78% | 23 | 33 | 70% | KV1 | 74% | 79% | 19 | 23 | 83% |
| Etaracizumab | V3 | 58% | 83% | 12 | 16 | 75% | KV3 | 62% | 69% | 13 | 25 | 52% |
| Talacotuzumab | V5 | 78% | 83% | 18 | 33 | 55% | KV4 | 73% | 73% | 11 | 16 | 69% |
| Rovalpituzumab | V1 | 67% | 67% | 21 | 30 | 70% | KV3 | 64% | 69% | 14 | 26 | 54% |
| Clazakizumab | V3 | 86% | 86% | 7 | 27 | 26% | KV1 | 75% | 75% | 4 | 22 | 18% |
| Ligelizumab | V1 | 64% | 64% | 11 | 21 | 52% | KV3 | 64% | 91% | 11 | 21 | 52% |
| Crizanlizumab | V1 | 64% | 64% | 11 | 23 | 48% | KV1 | 85% | 95% | 20 | 23 | 87% |
| Mogamulizumab | V3 | 67% | 67% | 6 | 15 | 40% | KV2 | 67% | 67% | 6 | 12 | 50% |
| Refanezumab | V7 | 87% | 87% | 15 | 17 | 88% | KV4 | 92% | 100% | 12 | 17 | 71% |
| Average |  | 68% | 77% |  |  | 59% |  | 77% | 85% |  |  | 58% |
| Median |  | 67% | 78% |  |  | 59% |  | 78% | 88% |  |  | 56% |

### Relationship of RF model scores with immunogenicity

The aim of humanization is to create a therapeutic that is safe and does not elicit an immune response. A strong predictive score for classification is not sufficient to produce a humanizer as it does not explicitly account for immunogenicity. The relationship of the model scores with observed immunogenic responses, as measured by the appearance of anti-drug antibodies (ADAs), was therefore investigated. The fraction of patients with observed immunogenic responses was obtained from FDA labels of approved antibody therapeutics and clinical studies of therapeutics still in clinical trials (outlined in Methods). There are limitations to this data: for example there are differences in patient demographic (age, physical conditions, illness), dosage levels and length of dosage of the therapeutic and if the treatment is in combination with other drugs. In addition, the murine therapeutics within the dataset are likely to be inherently biased towards lower levels of immunogenicity as they are approved therapeutics. SI section 3E shows the correlation between our model scores and the fraction of patients with observed immunogenic responses across 218 therapeutics. We found that higher model scores tend to relate to lower immunogenicity, although the correlation was weak with an R^2^ of 0.35. This correlation is significantly higher than the R^2^ of 0.18 observed in previous work (13).

We grouped the set of 481 therapeutics by their humanness scores (the output of the RF models). Figure 2 illustrates this categorization and demonstrates that high humanness scores are linked with low immunogenicity and vice versa. For example, over 90% of therapeutics that had both their VH and VL sequence above the YJS threshold exhibited low observed immunogenicity and only 1 sequence (0.7%) had high immunogenicity. In contrast, less than 50% of the therapeutics with scores below the YJS threshold had low immunogenicity.

**Fig 2.**
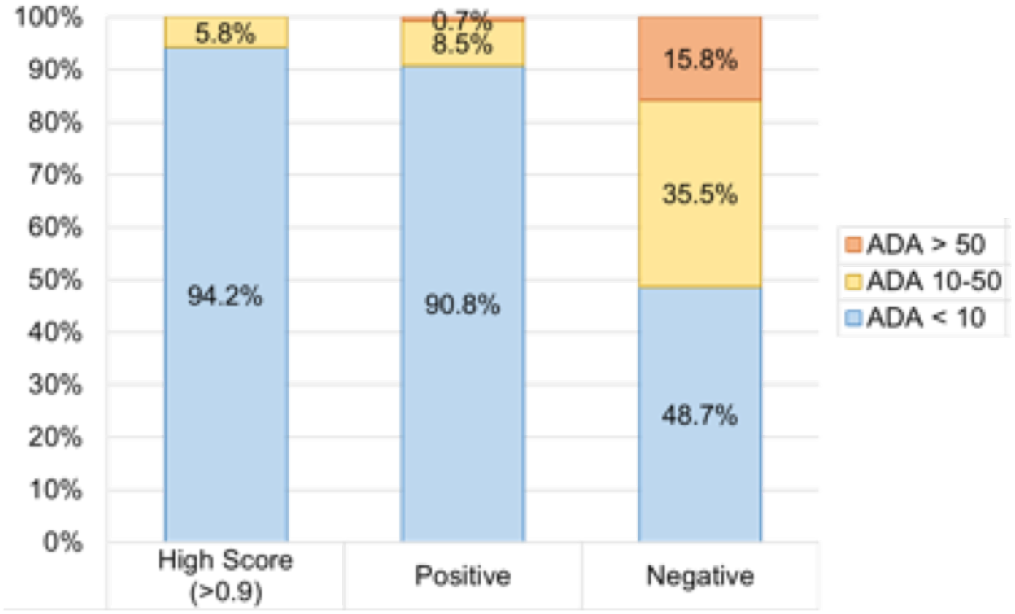
Relationship between the the humanness scores produced by our RF models and experimentally-determined immunogenicity. Therapeutics were split into three categories according to their humanness scores: above 0.9 (‘High Score’), above the YJS threshold for the relevant RF model (‘Positive’), and below the YJS threshold (‘Negative’). Both the VH and VL sequences have to be above the threshold to be classed as ‘Positive’. The immunogenicity of a therapeutic is also represented by three levels: over 50% of patients develop ADAs (orange), 10-50% of patients develop ADAs (yellow), and under 10% of patients develop ADAs (blue). Therapeutic sequences classified as human by our model tend to have low immunogenicity levels, while sequences classified as not human are more immunogenic.

**Fig 3.**
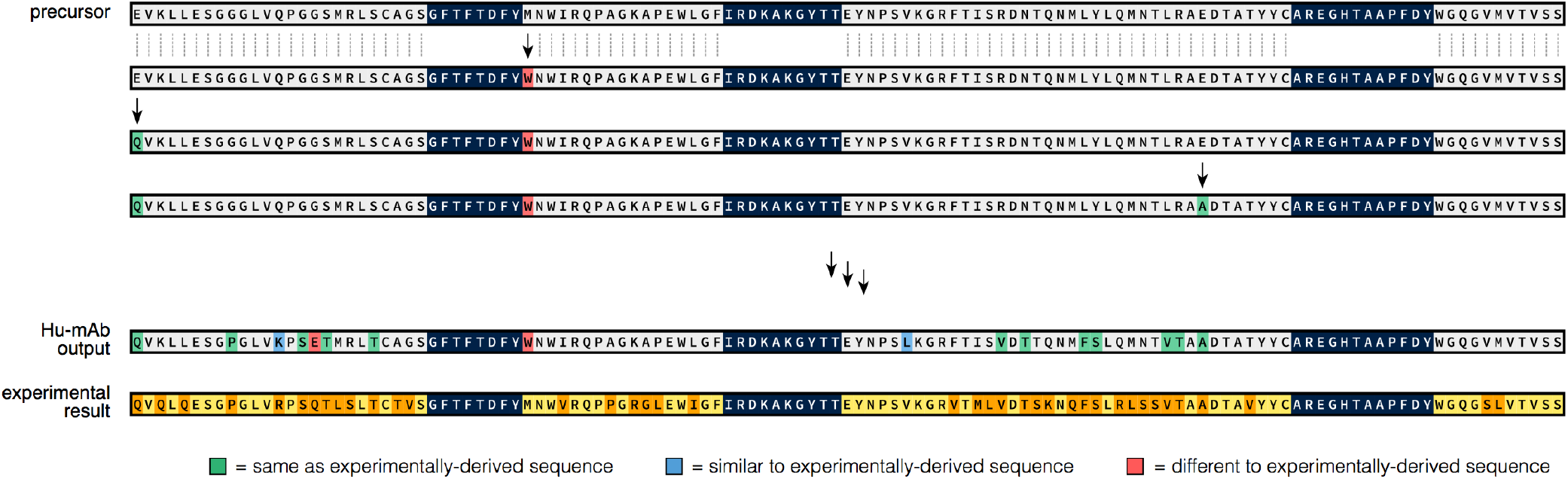
The Hu-mAb humanization procedure, demonstrated using the heavy chain sequence of the therapeutic Campath. The humanized sequence produced experimentally is shown at the bottom of the figure (conserved residues in yellow, mutated residues in orange). Starting with the unhumanized precursor sequence (top), Hu-mAb makes every possible mutation to the framework residues (grey) and selects the one that produces the largest increase in humanness score. CDR residues (dark blue) are not mutated to preserve binding. This procedure is performed iteratively until the humanness score reaches a given threshold. Mutations suggested by Hu-mAb are coloured depending on whether they are the same (green), similar (blue) or different (red) to mutations made experimentally. In this case, Hu-mAb suggested 16 mutations (compared to 39 from experiment), of which 14 were the same or similar to those derived experimentally.

**Fig 4.**
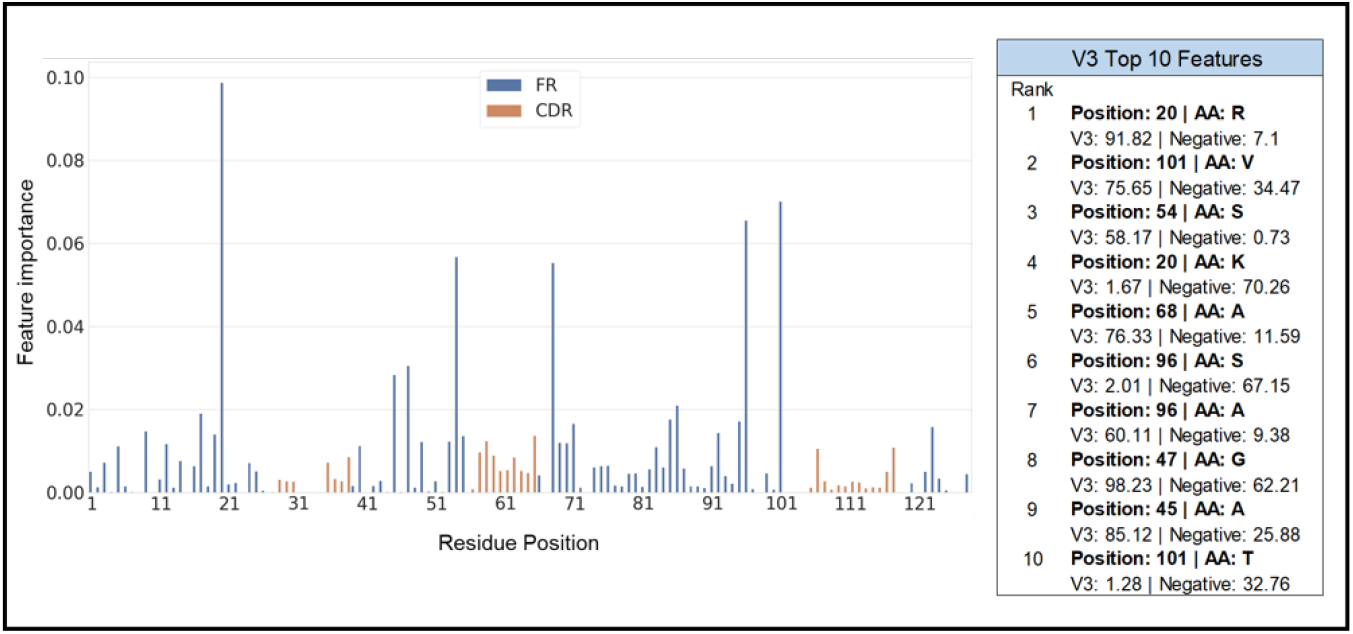
Feature importance of the VH V3 RF model and its top 10 features. The *x* axis consists of the residue positions in a sequential manner (left to right, IMGT numbering scheme). The inset table shows the top 10 features and the percentage frequency of the relevant amino acid type seen within the respective sets of sequences (V3 and negative, or non-human). The most important features likely determine the humanness of the sequence and are mainly located in the framework (FR) regions.

### Hu-mAb: a computational humanizer tool and its application to previously experimentally-humanized therapeutics

As high model scores were linked with lower levels of immunogenicity, we used the score to construct a computational humanization tool, Hu-mAb, that suggests optimal mutations that would increase the model score of the input sequence, therefore lowering immunogenicity. Residues in the CDRs are not mutated to maintain antigen-binding properties (described in Methods). The humanizer should ideally produce as few mutations as possible to reduce efficacy loss of the therapeutic. To investigate the similarity between mutations suggested by Hu-mAb and experimentally derived mutations, experimentally humanized sequences that demonstrated low immunogenicity and for which the precursor sequence was available were collected (for full details see SI sections 1E and 3F). The VH and VL sequence of each therapeutic was scored by each RF model, and the V gene identified by selecting the model that produced the highest score. The precursor sequence was used as the input sequence into the humanizer, along with its target humanness score (the score achieved by the experimentally-humanized sequence) and V gene type.

Table 1 compares the mutations made experimentally and those suggested by Hu-mAb for the precursor (unhumanized) sequences of 25 therapeutics. Each of these therapeutics displayed low immunogenicity in their experimentally-humanized forms. All precursor sequences were of murine, rat or rabbit origin and most had model scores close to 0 (see SI section 1E/3F for breakdown of scores and immunogenicity). Two therapeutics had precursor sequences scoring above their YJS threshold (VL only for Campath and both VH/VL for Clazakizumab). This is likely due to sequences of their species origin not being present in the training dataset of our models - neither VH/VL rabbit sequences (Clazakizumab) nor VL rat sequences (Campath) were present in the respective training datasets.

Hu-mAb consistently suggested fewer mutations than the number carried out experimentally on average, Hu-mAb suggested 59% and 58% of the experimental amount for the VH and VL sequences respectively. Of the mutations suggested by Hu-mAb, an average of 68% and 77% (for VH and VL sequences respectively) were also made experimentally (overlap ratio or OR). Including mutations to similar residue types (see SI section 3C for groupings) resulted in an average adjusted overlap ratio (AOR) of 77% and 85% for VH and VL respectively. This shows that the mutations suggested by Hu-mAb are very similar to those made experimentally. In contrast, a randomly humanized sequence would be expected to produce an average OR and AOR of *∼*2% and *∼*5% respectively (see SI section 3G). Hu-mAb is exploiting the information found in the antibody repertoires to more efficiently humanize therapeutic sequences.

### Hu-mAb protocol and RF model analysis

Since experimental humanization procedures often involve grafting of non-human CDRs onto a human framework, it is expected that the framework regions are more important than the hypervariable CDR regions for the classification of human and non-human sequences. Analysis of our RF models’ feature importance found that this is true; the key residues for discrimination are mostly found in the framework region 4. However, some CDR positions are utilized by the models for discrimination. The most important features (top 10) for each RF model are given in the Supplementary Information.

Analysis of our Hu-mAb protocol showed that identical mutations (i.e. mutations of position X to residue type Y) do not result in an identical increase in humanness score; the effect depends on the rest of the sequence. Moreover, we found that Hu-mAb occasionally made more than one mutation to the same position in the sequence over the course of the humanization procedure. These observations suggest that our RF models do not consider positions in the sequence independently, but rather they incorporate interactions between residues to more realistically evaluate humanness.

We have also analysed the characteristics of the mutations proposed by Hu-mAb and compared them to those made experimentally. In terms of residue types, the mutations proposed by Hu-mAb and experimentally were very similar (SI section 3H). Most commonly, mutations were from one hydrophobic residue to another (18% and 20% of all mutations made by Hu-mAb and through experiment, respectively). Least common were mutations involving cysteines (<1% for both Hu-mAb and experiment); importantly the conserved cysteines at IMGT positions 23 and 104 were never mutated, meaning structural viability is maintained (18).

The geometry of the antibody binding site is dependent on the orientation of the VH and VL, which is in turn affected by the residues present at the interface between the two domains. The proportion of mutations suggested by Hu-mAb to key VH-VL interface residues is slightly lower than the proportion made by experimental procedures (see SI section 3I), and the overlap ratio calculated for these residues is also higher than the average (74%/96% for VH/VL compared to an average across all mutations of 68%/77%). Since Hu-mAb also suggests fewer mutations on average (58-59% of the number made experimentally), the average number of interface mutations per sequence is around half that of experimental procedures (0.8 vs 1.6 for heavy chains, 0.8 vs 1.8 for light chains). A similar pattern was also observed for the Vernier zone - Hu-mAb proposed fewer mutations to these residues, which are thought to affect CDR conformations (19) (full details in SI). This means that the binding properties of the antibody are more likely to be preserved by using Hu-mAb.

## Discussion

We have developed a novel humanization tool (Hu-mAb) that can humanize both the VH and VL sequences of potential antibody therapeutics. The model is based on RF classifiers that have been trained on large-scale repertoire sequence data and demonstrate very high levels of accuracy in classification of antibodies by their origin. The humanness scores of the model exhibited a negative relationship with observed experimental immunogenicity. Therefore, sequences that have a higher humanness score are likely to have lower levels of immunogenicity.

Our model is worse at classifying non-human sequences of species that it has not been trained on (as seen with the rabbit precursor sequence of Clazakizumab). The non-human sequences within OAS are almost entirely of murine origin, and therefore Hu-mAb is mainly intended for use on murine precursor sequences. We intend to regularly train and update the RF models as new studies of non-human species are added to OAS, potentially widening its uses; however as most therapeutics of non-human origin are derived from murine sources, our RF models and humanizer Hu-mAb should already be applicable in many cases.

Experimental approaches to humanization are largely a trial-and-error process involving grafting of CDRs onto a completely human scaffold and if efficacy is lost, arbitrary back-mutations are made to attempt to restore it (8). Hu-mAb was constructed as a greedy algorithm and is optimized to select the mutations that provide the highest increase in humanness score, thus suggesting as few mutations as possible to reduce the likelihood of impacting the efficacy of the therapeutic. By utilizing RF classifiers that have only trained on a particular V gene type, the humanizer should produce a realistic sequence with a single V gene origin.

Hu-mAb is efficient and only proposes mutations to the key residues in the framework region responsible for humanness; it incrementally suggests additional mutations to reduce immunogenicity if necessary; and back-mutations can be suggested in a sequential and non-arbitrary manner (the mutation with the lowest impact on the humanness score). Compared to experimentally humanized therapeutics, Hu-mAb suggested 60% of the number of mutations, with a high similarity to those suggested experimentally (adjusted overlap ratio of 75-80%). Hu-mAb offers a promising alternative to experimental humanization approaches, allowing mutations to be made in a more systematic and efficient manner, and achieving similar results in a fraction of the time.

## Materials and Methods

### Preparation of OAS antibody sequence datasets

All IgG VH and VL sequences were downloaded from the OAS database (August 2020), totaling over 500 million sequences in the IMGT format. Human sequences were split by their V gene type - for example, V1 to V7 for VH sequences. Redundant sequences, sequences with cysteine errors (18) and sequences with missing framework 1 residues (residues preceding CDR1) were removed. The total dataset included over 65 million non-redundant sequences (SI section 1A).

### Training and testing the RF models

All models were trained using the scikit-learn Python module with default parameters unless stated otherwise. RF binary classifiers for each V gene type were trained with their respective set of V gene sequences and the entire set of negative sequences. For example, the VH V1 model was trained on all human VH V1 sequences (labelled as the positive class) and all VH negative sequences (labelled as the negative class). 80% of the dataset was used for training, 10% for validation and 10% for testing. Performance plateaued after 100-200 estimators and therefore each RF classifier was trained with 200 estimators. The performance of the RF models were assessed by determining its ability to correctly distinguish human sequences of a specific V gene type from those originating from other species. The validation set was utilized to set the classification threshold according to the value that maximizes the Youden’s J statistic (calculated as YJS = sensitivity + specificity - 1). It was found that the threshold that maximizes the YJS was very similar to the threshold that maximizes the Matthews correlation coefficient (see SI section 3B). This classification threshold was then used for calculating YJS values of the test set and for classification of therapeutic datasets. In addition, receiver operating characteristic (ROC) curves were generated and area under curve (AUC) scores for each model were calculated in order to assess performance.

### Training and testing the LSTM models

Identical training (excluding negative sequences), validation and test sets were used for the LSTM models. The method to construct the LSTM models followed that described in (14). As with the RF models, the validation set was used to set the classification threshold for the test dataset.

### VL kappa and lambda classifier

The RF model to classify whether a light chain sequence is of type kappa or lambda was trained on 25% of the total human VL dataset (12 million sequences). Testing of the model demonstrated perfect accuracy – it correctly classified every sequence as kappa or lambda within the entire VL dataset (both human and negative).

### Sequence alignments

All antibody sequences were aligned and numbered using the IMGT scheme with the AN-ARCI software (20).

### Therapeutic antibody dataset

All approved and phase 1-3 antibody therapeutics were obtained from Thera-SabDab (3) and were aligned and IMGT numbered by ANARCI (August 2020). Only mAbs with both a VH and VL sequence were included; this gave a set of 481 therapeutics (see SI section 1B). Each therapeutic has an International Nonproprietary Name (INN) assigned by the WHO (21). The INN infix preceding the suffix ‘-mab’ is determined by the origin of the therapeutic. Thus, the origin of each therapeutic was obtained from its source infix (SI sections 1C/2A). Therapeutics named in 2017 onwards no longer followed this nomenclature and their origins were obtained from the IMGT database for therapeutic monoclonal antibodies (IMGT/mab-DB) (22). The Supplementary Information contains lists of all 481 therapeutics and their origin, as well as a list of the 25 experimentally humanized therapeutic sequences and their precursors used to test Hu-mAb.

### ADA response levels of therapeutics

Anti-drug anti-body (ADA) responses of patients were obtained for 218 therapeutics from clinical papers using an identical approach to that described in (13). When multiple ADA levels are reported for the same therapeutic, the mean between the minimal and maximal reported value is used. The complete list of sequences together with observed immunogenicity levels can be found in the SI.

### Hu-mAb protocol

The input sequence, specific chain type (VH, kappa or lambda), V gene type, and target humanness score were used as inputs. To compare Hu-mAb to experimental mutations, for the therapeutic cases we set the Hu-mAb target score as the humanness score of the experimentally-humanized sequence. Every possible single site mutation within the framework region of the input sequence was made (SI section 2B). This generated a set of mutated sequences which were then scored by the relevant RF model. The humanness scores of the mutated sequences were ranked and the top scoring sequence was selected. This process was repeated with the newly selected sequence until the target humanness score was achieved.

## Supporting information

SI - Full Hu-mAb Results

SI - Therapeutic Sequences

SI - Therapeutic ADA values

Supplementary information (main)

