## Supplementary material for "Humanization of antibodies using a machine learning approach on large-scale repertoire data": SI - Full Hu-mAb Results

**Experimentally-humanised sequences vs Hu-mAb output for all therapeutics**

Experimental sequences:
- = not mutated
- = mutated

Hu-mAb sequences:
- = not mutated
- = identical to experimentally-humanised sequence
- = similar to experimentally-humanised sequence
- = different to experimentally-humanised sequence

## CD28

Heavy Chain

| | |
|---|---|
| # experimental mutations: | 33 |
| # humab mutations: | 19 |
| mutation ratio: | 58% |
| overlap ratio: | 63% |
| adjusted OR: | 79% |

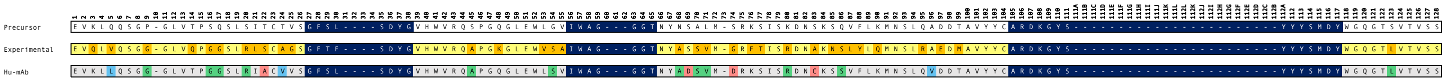

Light Chain

| | |
|---|---|
| # experimental mutations: | 19 |
| # humab mutations: | 11 |
| mutation ratio: | 58% |
| overlap ratio: | 64% |
| adjusted OR: | 73% |

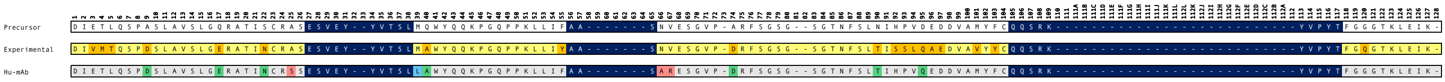

#### Campath

Heavy Chain

| | |
|---|---|
| # experimental mutations: | 39 |
| # humab mutations: | 16 |
| mutation ratio: | 41% |
| overlap ratio: | 75% |
| adjusted OR: | 88% |

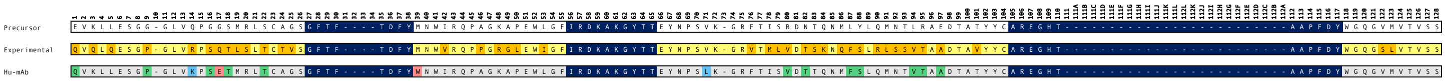

Light Chain

| | |
|---|---|
| # experimental mutations: | 14 |
| # humab mutations: | 3 |
| mutation ratio: | 21% |
| overlap ratio: | 67% |
| adjusted OR: | 67% |

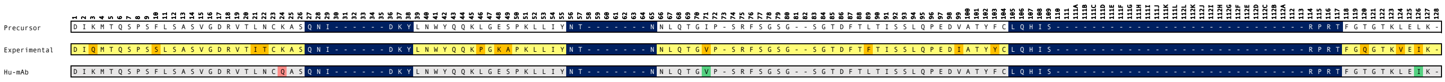

#### Bevacizumab

Heavy Chain

| | |
|---|---|
| # experimental mutations: | 25 |
| # humab mutations: | 14 |
| mutation ratio: | 56% |
| overlap ratio: | 50% |
| adjusted OR: | 57% |

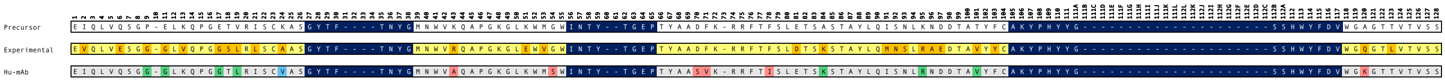

Light Chain

| | |
|---|---|
| # experimental mutations: | 16 |
| # humab mutations: | 9 |
| mutation ratio: | 56% |
| overlap ratio: | 89% |
| adjusted OR: | 100% |

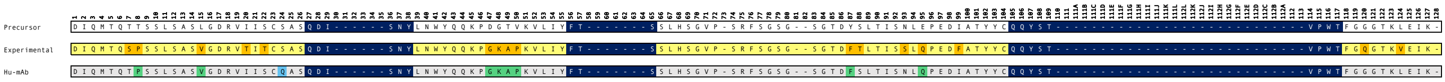

#### Herceptin

Heavy Chain

| | |
|---|---|
| # experimental mutations: | 32 |
| # humab mutations: | 27 |
| mutation ratio: | 84% |
| overlap ratio: | 59% |
| adjusted OR: | 78% |

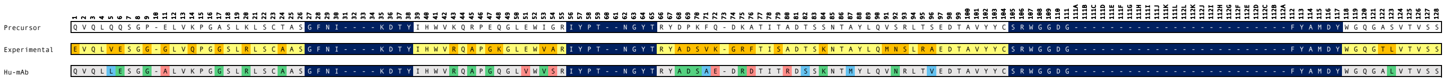

Light Chain

| | |
|---|---|
| # experimental mutations: | 22 |
| # humab mutations: | 8 |
| mutation ratio: | 36% |
| overlap ratio: | 88% |
| adjusted OR: | 88% |

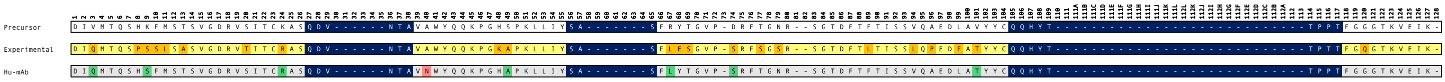

#### Omalizumab

Heavy Chain

| | |
|---|---|
| # experimental mutations: | 34 |
| # humab mutations: | 21 |
| mutation ratio: | 62% |
| overlap ratio: | 62% |
| adjusted OR: | 76% |

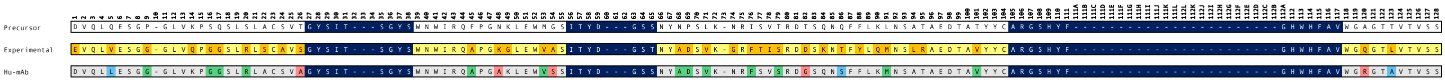

Light Chain

| | |
|---|---|
| # experimental mutations: | 25 |
| # humab mutations: | 19 |
| mutation ratio: | 76% |
| overlap ratio: | 89% |
| adjusted OR: | 95% |

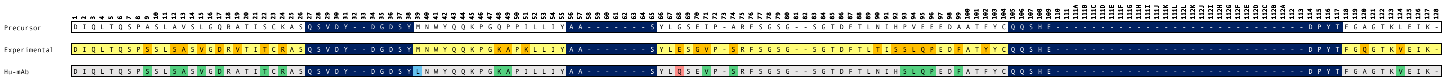

#### Eculizumab

##### Heavy Chain

| | |
|---|---|
| # experimental mutations: | 23 |
| # humab mutations: | 15 |
| mutation ratio: | 65% |
| overlap ratio: | 73% |
| adjusted OR: | 73% |

| | Sequence |
|---|---|
| Precursor | QVQLQQSGA-ELMKPGASVKMSCKATGYIF----SNYWIQWIKQRPGHGLEWIGEILPG--SGSTEYTENFK-DKAAFTADTSSNTAYMQLSSLTSEDSAVYYCARYFFGSS----------------------PNWYFDVWGAGTTVTVSS |
| Experimental | QVQLVQSGA-EVKKPGASVKVSCKASGYIF----SNYWIQWVRQAPGQGLEWMGEILPG--SGSTEYTENFK-DRVTMTRDTSTSTVYMELSSLRSEDTAVYYCARYFFGSS----------------------PNWYFDVWGQGTLVTVSS |
| Hu-mAb | QVQLVQSGA-EVKKPGASVKVSCKATGYIF----SNYWIQWIRQAPGHGLEWMGWILPG--SGSTEYTENFQ-DRVAFTADTSINTAYMELSSLTSEDTAVYYCARYFFGSS----------------------PNWYFDVWGCGTTVTVSS |

##### Light Chain

| | |
|---|---|
| # experimental mutations: | 20 |
| # humab mutations: | 12 |
| mutation ratio: | 60% |
| overlap ratio: | 83% |
| adjusted OR: | 83% |

| | Sequence |
|---|---|
| Precursor | DIQMTQSPASLSASVGETVTITCGASENI------YGALNWYQRKQGKSPQLLIYGA-------TNLADGMS-SRFSGSG--SGRQYYLKISSLHPDDVATYYCQNVLN----------------------------TPLTFGAGTKLELK- |
| Experimental | DIQMTQSPSSLSASVGDRVTITCGASENI------YGALNWYQQKPGKAPKLLIYGA-------TNLADGVP-SRFSGSG--SGTDFTLTISSLQPEDFATYYCQNVLN----------------------------TPLTFGQGTKVEIK- |
| Hu-mAb | DIQMTQSPSSLSASVGDRVTITCRASENI------YGALNWYQRKPGKAPQLLIYGA-------TNLQDGVS-SRFSGSG--SGTQFYLTISSLHPDDVATYYCQNVLN----------------------------TPLTFGQGTKLELK- |

#### Tocilizumab

##### Heavy Chain

| | |
|---|---|
| # experimental mutations: | 23 |
| # humab mutations: | 14 |
| mutation ratio: | 61% |
| overlap ratio: | 64% |
| adjusted OR: | 86% |

| | Sequence |
|---|---|
| Precursor | DVQLQESGP-VLVKPSQSLSLTCTVTGYSIT---SDHAWSWIRQFPGNKLEWMGYISYS---GITTYNPSLK-SRISITRDTSKNQFFLQLNSVTTGDTSTYYCARSLAR-------------------------TTAMDYWGQGTSVTVSS |
| Experimental | QVQLQESGP-GLVRPSQTLSLTCTVSGYSIT---SDHAWSWVRQPPGRGLEWIGYISYS---GITTYNPSLK-SRVTMLRDTSKNQFSLRLSSVTAADTAVYYCARSLAR-------------------------TTAMDYWGQGSLVTVSS |
| Hu-mAb | QLQLQESGP-VLVKPSETLSLTCTVSGYSIT---SDHAWSWIRQPPGKGLEWIGYISYS---GITTYNPSLK-SRASITVDTSKNQFSLQLNSVTAGDTSVYYCARSLAR-------------------------TTAMDYWGQGTSVTVSS |

##### Light Chain

| | |
|---|---|
| # experimental mutations: | 19 |
| # humab mutations: | 9 |
| mutation ratio: | 47% |
| overlap ratio: | 78% |
| adjusted OR: | 89% |

| | Sequence |
|---|---|
| Precursor | DIQMTQTTSSLSASLGDRVTISCRASQDI------SSYLNWYQQKPDGTIKLLIYYT-------SRLHSGVP-SRFSGSG--SGTDYSLTINNLEQEDIATYFCQQGNT----------------------------LPYTFGGGTKLEIN- |
| Experimental | DIQMTQSPSSLSASVGDRVTITCRASQDI------SSYLNWYQQKPGKAPKLLIYYT-------SRLHSGVP-SRFSGSG--SGTDFTFTISSLQPEDIATYYCQQGNT----------------------------LPYTFGQGTKVEIK- |
| Hu-mAb | DIQMTQTTSSLSASVGDRVTISCRASQDI------SSYLNWYQQKPDKAPKLLIYYT-------SRLQSGVP-SRFSGSG--SGTDFSLTINNLQQEDIATYFCQQGNT----------------------------LPYTFGQGTRLEIN- |

#### Pembrolizumab

##### Heavy Chain

| | |
|---|---|
| # experimental mutations: | 23 |
| # humab mutations: | 11 |
| mutation ratio: | 48% |
| overlap ratio: | 73% |
| adjusted OR: | 73% |

| | Sequence |
|---|---|
| Precursor | QVQLQQPGA-ELVKPGTSVKLSCKASGYTF----TNYYMYWVKQRPGQGLEWIGGINPS--NGGTNFNEKFK-NKATLTVDSSSSTTYMQLSSLTSEDSAVYYCTRRDYRF------------------------DMGFDYWGQGTTLTVSS |
| Experimental | QVQLVQSGV-EVKKPGASVKVSCKASGYTF----TNYYMYWVRQAPGQGLEWMGGINPS--NGGTNFNEKFK-NRVTLTTDSSTTTAYMELKSLQFDDTAVYYCARRDYRF------------------------DMGFDYWGQGTTVTVSS |
| Hu-mAb | QVQLVQPGA-EVKKPGTSVKLSCKASGYTF----TNYYMYWVRQAPGQGLEWMGGINPS--NGGTNFAEKFQ-NKVTLTVDSSISTTYMELSSLTSEDSAVYYCTRRDYRF------------------------DMGFDYWGQGTTLTVSS |

##### Light Chain

| | |
|---|---|
| # experimental mutations: | 20 |
| # humab mutations: | 12 |
| mutation ratio: | 60% |
| overlap ratio: | 75% |
| adjusted OR: | 75% |

| | Sequence |
|---|---|
| Precursor | DIVLTQSPASLAVSLGQRAAISCRASKGVST--SGYSYLHWYQQKPGQSPKLLIYLA-------SYLESGVP-ARFSGSG--SGTDFTLNIHPVEEEDAATYYCQHSRD----------------------------LPLTFGTGTKLELK- |
| Experimental | EIVLTQSPATLSLSPGERATLSCRASKGVST--SGYSYLHWYQQKPGQAPRLLIYLA-------SYLESGVP-ARFSGSG--SGTDFTLTISSLEPEDFAVYYCQHSRD----------------------------LPLTFGGGTKVEIK- |
| Hu-mAb | EIVLTQSPATLAVSPGERAALSCRASKGVST--SGYSYLAWYQQKPGQSPRLLIYLA-------SYWASGVP-ARFSGSG--SGTDFTLNIHSLEEEDFATYYCQHSRD----------------------------LPLTFGTGTKLELK- |

#### Pertuzumab

##### Heavy Chain

| | |
|---|---|
| # experimental mutations: | 32 |
| # humab mutations: | 19 |
| mutation ratio: | 59% |
| overlap ratio: | 68% |
| adjusted OR: | 79% |

| | Sequence |
|---|---|
| Precursor | EVQLQQSGP-ELVKPGTSVKISCKASGFTF----TDYTMDWVKQSHGKSLEWIGDVNPN--SGGSIYNQRFK-GKASLTVDRSSRIVYMELRSLTFEDTAVYYCARNLGP-------------------------SFYFDYWGQGTTLTVSS |
| Experimental | EVQLVESGG-GLVQPGGSLRLSCAASGFTF----TDYTMDWVRQAPGKGLEWVADVNPN--SGGSIYNQRFK-GRFTLSVDRSKNTLYLQMNSLRAEDTAVYYCARNLGP-------------------------SFYFDYWGQGTLVTVSS |
| Hu-mAb | EVQLLQSGG-TLVKPGGSLRISCVASGFTF----TDYTMDWVRQAHGKSLEWVSDVNPN--SGGSIYAQRGK-GRFSLSVDRSKRIVYMEMRSLTFEDTAVYYCARNLGP-------------------------SFYFDYWGQGTTVTVSS |

##### Light Chain

| | |
|---|---|
| # experimental mutations: | 20 |
| # humab mutations: | 10 |
| mutation ratio: | 50% |
| overlap ratio: | 80% |
| adjusted OR: | 90% |

| | Sequence |
|---|---|
| Precursor | DTVMTQSHKIMSTSVGDRVSITCKASQDV------SIGVAWYQQRPGQSPKLLIYSA-------SYRYTGVP-DRFTGSG--SGTDFTFTISSVQAEDLAVYYCQQYYI----------------------------YPYTFGGGTKLEIK- |
| Experimental | DIQMTQSPSSLSASVGDRVTITCKASQDV------SIGVAWYQQKPGKAPKLLIYSA-------SYRYTGVP-SRFSGSG--SGTDFTLTISSLQPEDFATYYCQQYYI----------------------------YPYTFGQGTKVEIK- |
| Hu-mAb | DTQMTQSHSIMSASVGDRVSITCRASQDV------SIGVAWYQQRPGQAPKLLIYSA-------SYLYTGVP-SRFTGSG--SGTDFTFTISSLQPEDLATYYCQQYYI----------------------------YPYTFGGGTKLEIK- |

#### Ixekizumab

##### Heavy Chain

| | |
|---|---|
| # experimental mutations: | 29 |
| # humab mutations: | 12 |
| mutation ratio: | 41% |
| overlap ratio: | 75% |
| adjusted OR: | 75% |

| | Sequence |
|---|---|
| Precursor | QVQLQQSRP-ELVKPGASVKISCKASGYSF----TDYNMNWVKQSNGKSLEWIGVINPN--YGTTDYNQRFK-GKATLTVDQSSRTAYMQLNSLTSEDSAVYYCVIYDYA-------------------------TGTGGYWGQGSPLTVSS |
| Experimental | QVQLVQSGA-EVKKPGSSVKVSCKASGYSF----TDYHIHWVRQAPGQGLEWMGVINPM--YGTTDYNQRFK-GRVTITADESTSTAYMELSSLRSEDTAVYYCARYDYF-------------------------TGTGVYWGQGTLVTVSS |
| Hu-mAb | QVQLVQSRA-EVKKPGASVKISCKASGYSF----TDYNMNWVRQANGKSLEWMGVINPN--YGTTDYAQRFQ-GKVTLTVDQSIRTAYMELNSLTSEDSAVYYCVIYDYA-------------------------TGTGGYWGQGSPLTVSS |

##### Light Chain

| | |
|---|---|
| # experimental mutations: | 12 |
| # humab mutations: | 9 |
| mutation ratio: | 75% |
| overlap ratio: | 78% |
| adjusted OR: | 100% |

| | Sequence |
|---|---|
| Precursor | DVVLTQTPLSLPVSLGDQASISCRSSQSLVHS-NGNTYLHWYLQKPGQSPKLLIYKV-------SNRFSGVP-DRFSGSG--SGTDFTLKISRVEAEDLGVYFCSQSTH----------------------------VPFTFGSGTKLEIK- |
| Experimental | DIVMTQTPLSLSVTPGQPASISCRSSRSLVHS-RGNTYLHWYLQKPGQSPQLLIYKV-------SNRFIGVP-DRFSGSG--SGTDFTLKISRVEAEDVGVYYCSQSTH----------------------------LPFTFGQGTKLEIK- |
| Hu-mAb | DVVLTQSPLSLPVTLGQPASISCRSSQSLVHS-NGNTYLHWYLQRPGQSPQLLIYKV-------SNRFSGVP-DRFSGSG--SGTDFTLKISRVEAEDVGVYYCSQSTH----------------------------VPFTFGQGTKLEIK- |

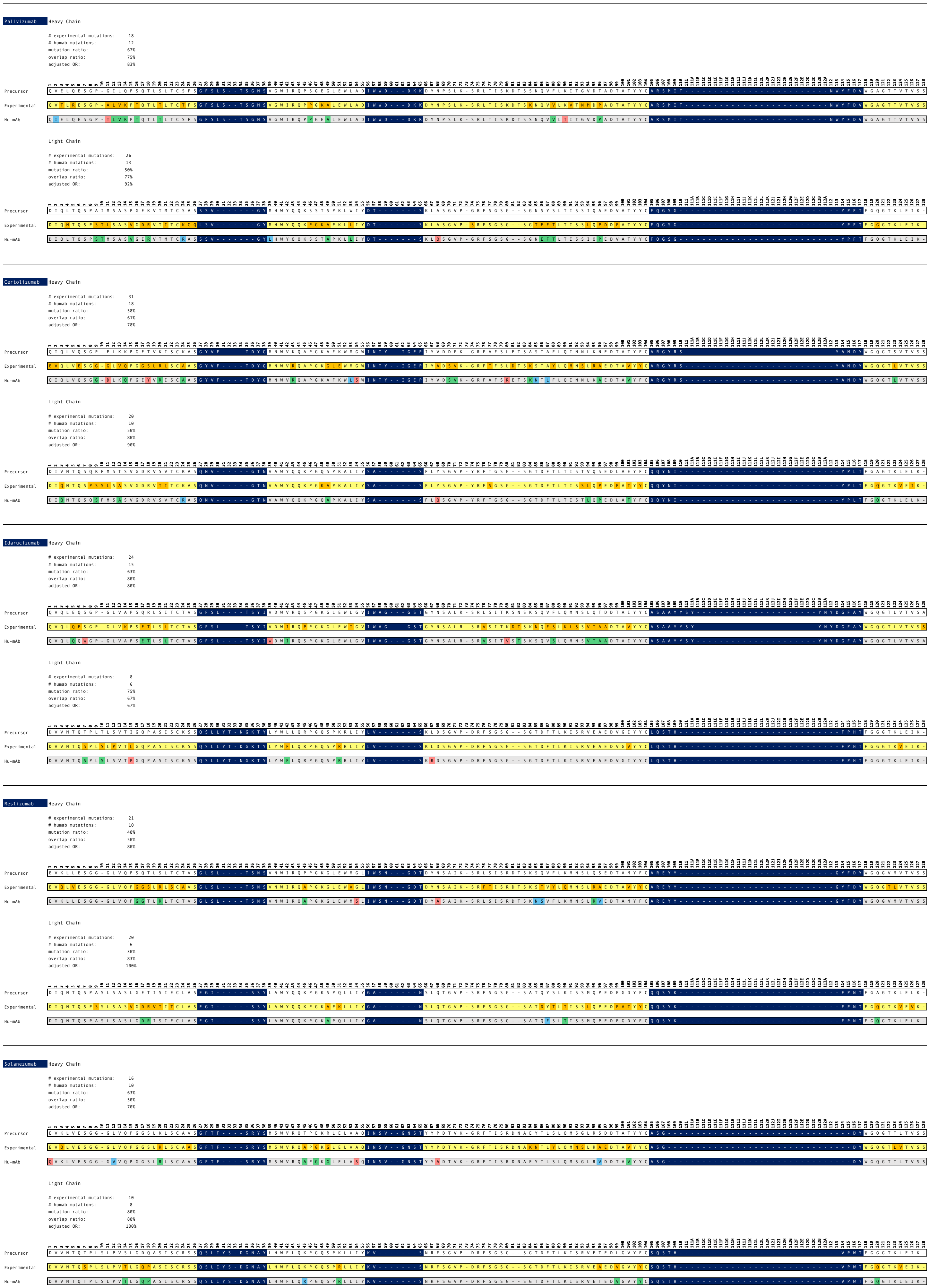

Palivizumab
Heavy Chain
### experimental mutations: 18
### humab mutations: 12
mutation ratio: 67%
overlap ratio: 75%
adjusted OR: 83%
Precursor
Experimental
Hu-mAb
Light Chain
### experimental mutations: 26
### humab mutations: 13
mutation ratio: 50%
overlap ratio: 77%
adjusted OR: 92%
Precursor
Experimental
Hu-mAb
Certolizumab
Heavy Chain
### experimental mutations: 31
### humab mutations: 18
mutation ratio: 58%
overlap ratio: 61%
adjusted OR: 78%
Precursor
Experimental
Hu-mAb
Light Chain
### experimental mutations: 20
### humab mutations: 10
mutation ratio: 50%
overlap ratio: 80%
adjusted OR: 90%
Precursor
Experimental
Hu-mAb
Idarucizumab
Heavy Chain
### experimental mutations: 24
### humab mutations: 15
mutation ratio: 63%
overlap ratio: 80%
adjusted OR: 80%
Precursor
Experimental
Hu-mAb
Light Chain
### experimental mutations: 8
### humab mutations: 6
mutation ratio: 75%
overlap ratio: 67%
adjusted OR: 67%
Precursor
Experimental
Hu-mAb
Reslizumab
Heavy Chain
### experimental mutations: 21
### humab mutations: 10
mutation ratio: 48%
overlap ratio: 50%
adjusted OR: 80%
Precursor
Experimental
Hu-mAb
Light Chain
### experimental mutations: 20
### humab mutations: 6
mutation ratio: 30%
overlap ratio: 83%
adjusted OR: 100%
Precursor
Experimental
Hu-mAb
Solanezumab
Heavy Chain
### experimental mutations: 16
### humab mutations: 10
mutation ratio: 63%
overlap ratio: 50%
adjusted OR: 70%
Precursor
Experimental
Hu-mAb
Light Chain
### experimental mutations: 10
### humab mutations: 8
mutation ratio: 80%
overlap ratio: 88%
adjusted OR: 100%
Precursor
Experimental
Hu-mAb

#### Lorvotuzumab

##### Heavy Chain

| | |
|---|---|
| # experimental mutations: | 13 |
| # humab mutations: | 10 |
| mutation ratio: | 77% |
| overlap ratio: | 90% |
| adjusted OR: | 90% |

| | |
|---|---|
| Precursor | DVQLVESGG-GLVQPGGSRKLSCAASGFTF----SSFGMHWVRQAPEKGLEWVAYISSG--SFTIYHADTVK-GRFTISRDNPKNTLFLQMTSLRAEDTAHYYCARMRKG--------------------------YAMDYWGQGTTVTVSS |
| Experimental | QVQLVESGG-GVVQPGRSLRLSCAASGFTF----SSFGMHWVRQAPGKGLEWVAYISSG--SFTIYYADSVK-GRFTISRDNSKNTLYLQMNSLRAEDTAVYYCARMRKG--------------------------YAMDYWGQGTLVTVSS |
| Hu-mAb | QVQLVESGG-GVVQPGGSLRLSCAASGFTF----SSFGMHWVRQAPGKGLEWVSYISSG--SFTIYHADSVK-GRFTISRDNSKNTLFLQMNSLRAEDTAHYYCARMRKG--------------------------YAMDYWGQGTLVTVSS |

##### Light Chain

| | |
|---|---|
| # experimental mutations: | 13 |
| # humab mutations: | 11 |
| mutation ratio: | 85% |
| overlap ratio: | 82% |
| adjusted OR: | 82% |

| | |
|---|---|
| Precursor | DVLMTQTPLSLPVSLGDQASISCRSSQIIIHS-DGNTYLEWFLQKPGQSPKLLIYKV-------SNRFSGVP-DRFSGSG--SGTDFTLMISRVEAEDLGVYYCFQGSH----------------------------VPHTFGGGTKLEIK- |
| Experimental | DVVMTQSPLSLPVTLGQPASISCRSSQIIIHS-DGNTYLEWFQQRPGQSPRRLIYKV-------SNRFSGVP-DRFSGSG--SGTDFTLKISRVEAEDVGVYYCFQGSH----------------------------VPHTFGQGTKVEIK- |
| Hu-mAb | DVLMTQSPLSLPVTLGQPASISCRSSQIIIHS-DGNTYLYWFQQRPGQSPQRLIYKV-------SNRFSGVP-DRFSGSG--SGTDFTLMISRVEAEDVGVYYCFQGSH----------------------------VPHTFGQGTKLEIK- |

#### Pinatuzumab

##### Heavy Chain

| | |
|---|---|
| # experimental mutations: | 33 |
| # humab mutations: | 23 |
| mutation ratio: | 70% |
| overlap ratio: | 61% |
| adjusted OR: | 78% |

| | |
|---|---|
| Precursor | QVQLQQSGP-ELVKPGASVKISCKASGYEF----SRSWMNWVKQRPGQGREWIGRIYPG--DGDTNYSGKFK-GKATLTADKSSSTAYMQLSSLTSVDSAVYFCARDGSSW------------------------DWYFDVWGAGTTVTVSS |
| Experimental | EVQLVESGG-GLVQPGGSLRLSCAASGYEF----SRSWMNWVRQAPGKGLEWVGRIYPG--DGDTNYSGKFK-GRFTISADTSKNTAYLQMNSLRAEDTAVYYCARDGSSW------------------------DWYFDVWGQGTLVTVSS |
| Hu-mAb | QVQLVQSGG-ELIQPGGSLRISCVASGYEF----SRSWMNWVRQAPGQGREWVSRIYPG--DGDTNYSGSFK-GAFTLSRDKSKSTLYMQMSSLTVVDTAVYFCARDGSSW------------------------DWYFDVWGCGTTVTVSS |

##### Light Chain

| | |
|---|---|
| # experimental mutations: | 23 |
| # humab mutations: | 19 |
| mutation ratio: | 83% |
| overlap ratio: | 74% |
| adjusted OR: | 79% |

| | |
|---|---|
| Precursor | DILMTQTPLSLPVSLGDQASISCRSSQSIVHS-NGNTFLEWYLQKPGQSPKLLIYKV-------SNRFSGVP-DRFSGSG--SGTDFTLKISRVEAEDLGVYYCFQGSQ----------------------------FPYTFGGGTKVEIK- |
| Experimental | DIQMTQSPSSLSASVGDRVTITCRSSQSIVHS-VGNTFLEWYQQKPGKAPKLLIYKV-------SNRFSGVP-SRFSGSG--SGTDFTLTISSLQPEDFATYYCFQGSQ----------------------------FPYTFGQGTKVEIK- |
| Hu-mAb | AIQMTQSPSTLPASVGDRASISCRASQSIVHS-NGNTFLEWYLQKPGKAPKLLIYKV-------SNLQSGVP-SRFSGSG--SGTDFTLTISRLQPEDLGTYYCFQGSQ----------------------------FPYTFGGGTKVEIK- |

#### Etaracizumab

##### Heavy Chain

| | |
|---|---|
| # experimental mutations: | 16 |
| # humab mutations: | 12 |
| mutation ratio: | 75% |
| overlap ratio: | 58% |
| adjusted OR: | 83% |

| | |
|---|---|
| Precursor | EVQLEESGG-GLVKPGGSLKLSCAASGFAF----SSYDMSWVRQIPEKRLEWVAKVSSG--GGSTYYLDTVQ-GRFTISRDNAKNTLYLQMSSLNSEDTAMYYCARHNY---------------------------GSFAYWGQGTLVTVSA |
| Experimental | QVQLVESGG-GVVQPGRSLRLSCAASGFTF----SSYDMSWVRQAPGKGLEWVAKVSSG--GGSTYYLDTVQ-GRFTISRDNSKNTLYLQMNSLRAEDTAVYYCARHLH---------------------------GSFASWGQGTTVTVSS |
| Hu-mAb | QVQLLESGG-GLVKPGGSLRLSCAASGFAF----SSYDMSWVRQIPGKGLEWVSKVSSG--GGSTYYADFVQ-GRFTISRDNAKNTLYLQMNSLNVEDTAVYYCARHNY---------------------------GSFAYWGQGTLVTVSS |

##### Light Chain

| | |
|---|---|
| # experimental mutations: | 25 |
| # humab mutations: | 13 |
| mutation ratio: | 52% |
| overlap ratio: | 62% |
| adjusted OR: | 69% |

| | |
|---|---|
| Precursor | ELVMTQTPATLSVTPGDSVSLSCRASQSI------SNHLHWYQQKSHESPRLLIKYA-------SQSISGIP-SRFSGSG--SGTDFTLSINSVETEDFGMYFCQQSNS----------------------------WPHTFGGGTKLEIK- |
| Experimental | EIVLTQSPATLSLSPGERATLSCQASQSI------SNFLHWYQQRPGQAPRLLIRYR-------SQSISGIP-ARFSGSG--SGTDFTLTISSLEPEDFAVYYCQQSGS----------------------------WPLTFGGGTKVEIK- |
| Hu-mAb | ELVMTQTPATLSVSPGDSASLSCRASQSI------SNHLAWYQQKSGQSPRLLIYYA-------SFRASGIP-ARFSGSG--SGTDFTLSINSLETEDFAMYYCQQSNS----------------------------WPHTFGGGTKLEIK- |

#### Talacoluzumab

##### Heavy Chain

| | |
|---|---|
| # experimental mutations: | 33 |
| # humab mutations: | 18 |
| mutation ratio: | 55% |
| overlap ratio: | 78% |
| adjusted OR: | 83% |

| | |
|---|---|
| Precursor | EVQLQQSGP-ELVKPGASVKMSCKASGYTF----TDYYMKWVKQSHGKSLEWIGDIIPS--NGATFYNQKFK-GKATLTVDRSSSTAYMHLNSLTSEDSAVYYCTRSHLLR------------------------ASWFAYWGQGTLVTVSA |
| Experimental | EVQLVQSGA-EVKKPGESLKISCKGSGYSF----TDYYMKWARQMPGKGLEWMGDIIPS--NGATFYNQKFK-GQVTISADKSISTTYLQWSSLKASDTAMYYCARSHLLR------------------------ASWFAYWGQGTMVTVSS |
| Hu-mAb | EVQLVQSGA-EVKKPGESLKMSCKASGYTF----TDYYIGWVRQSHGKSLEWIGDIIPS--NGATFYNQSFQ-GQVTLSVDRSISTAYMHWNSLTASDSAVYYCTRSHLLR------------------------ASWFAYWGQGTLVTVSA |

##### Light Chain

| | |
|---|---|
| # experimental mutations: | 16 |
| # humab mutations: | 11 |
| mutation ratio: | 69% |
| overlap ratio: | 73% |
| adjusted OR: | 73% |

| | |
|---|---|
| Precursor | DFVMTQSPSSLTVTAGEKVTMSCKSSQSLLNSGNQKNYLTWYLQKPGQPPKLLIYWA-------STRESGVP-DRFTGSG--SGTDFTLTISSVQAEDLAVYYCQNDYS----------------------------YPYTFGGGTKLEIK- |
| Experimental | DIVMTQSPDSLAVSLGERATINCESSQSLLNSGNQKNYLTWYQQKPGQPPKPLIYWA-------STRESGVP-DRFSGSG--SGTDFTLTISSLQAEDVAVYYCQNDYS----------------------------YPYTFGQGTKLEIK- |
| Hu-mAb | DFVMTQSPDSLTVALGERATFNCKSSQSLLNSGNQKNYLTWYLQKPGQPPKLLIYWA-------STRESGVP-DRFSGSG--SGTDFTLTISSLQPEDVAVYYCQNDYS----------------------------YPYTFGGGTKLEIK- |

#### Rovalpituzumab

##### Heavy Chain

| | |
|---|---|
| # experimental mutations: | 30 |
| # humab mutations: | 21 |
| mutation ratio: | 70% |
| overlap ratio: | 67% |
| adjusted OR: | 67% |

| | |
|---|---|
| Precursor | QIQLVQSGP-ELKKPGETVKISCKASGYTF----TNYGMNWVKQAPGKGLKWMAWINTY--TGEPTYADDFK-GRFAFSLETSASTASLQIINLKNEDTATYFCARIGDS--------------------------SPSDYWGQGTTLTVSS |
| Experimental | QVQLVQSGA-EVKKPGASVKVSCKASGYTF----TNYGMNWVRQAPGQGLEWMGWINTY--TGEPTYADDFK-GRVTMTTDTSTSTAYMELRSLRSDDTAVYYCARIGDS--------------------------SPSDYWGQGTLVTVSS |
| Hu-mAb | QVQLVQSGA-EVKKPGASVKVSCKASGYTF----TNYGMNWVRQAPGQGLKWMAWINTY--TGEPKYAQDFQ-GQVAMTAETSISTASMEIINLKAEDTAVYFCARIGDS--------------------------SPSDYWGQGTTLTVSS |

##### Light Chain

| | |
|---|---|
| # experimental mutations: | 26 |
| # humab mutations: | 14 |
| mutation ratio: | 54% |
| overlap ratio: | 64% |
| adjusted OR: | 79% |

| | |
|---|---|
| Precursor | SIVMTQTPKFLLVSAGDRVTITCKASQSV------SNDVVWYQQKPGQSPKLLIYYA-------SNRYTGVP-DRFAGSG--YGTDFSFTISTVQAEDLAVYFCQQDYT----------------------------SPWTFGGGTKLEIR- |
| Experimental | EIVMTQSPATLSVSPGERATLSCKASQSV------SNDVVWYQQKPGQAPRLLIYYA-------SNRYTGIP-ARFSGSG--SGTEFTLTISSLQSEDFAVYYCQQDYT----------------------------SPWTFGQGTKLEIK- |
| Hu-mAb | EIVLTQTPGTLSVSAGDRATLTCKASQSV------SNDAVWYQQKPGQSPRLLIYYA-------SNRATGVP-DRFAGSG--YGTDFSLTISTLEAEDFAVYFCQQDYT----------------------------SPWTFGGGTKLEIR- |

Numbering (printed above every sequence block): 1 2 3 4 5 6 7 8 9 10 11 12 13 14 15 16 17 18 19 20 21 22 23 24 25 26 27 28 29 30 31 32 33 34 35 36 37 38 39 40 41 42 43 44 45 46 47 48 49 50 51 52 53 54 55 56 57 58 59 60 61 62 63 64 65 66 67 68 69 70 71 72 73 74 75 76 77 78 79 80 81 82 83 84 85 86 87 88 89 90 91 92 93 94 95 96 97 98 99 100 101 102 103 104 105 106 107 108 109 110 111 111A 111B 111C 111D 111E 111F 111G 111H 111I 111J 111K 111L 112L 112K 112J 112I 112H 112G 112F 112E 112D 112C 112B 112A 112 113 114 115 116 117 118 119 120 121 122 123 124 125 126 127 128

**Clazakizumab** Heavy Chain

### experimental mutations: 27
### humab mutations: 7
mutation ratio: 26%
overlap ratio: 86%
adjusted OR: 86%

| | |
|---|---|
| Precursor | Q-SLEESGG-RLVTPGTPLTLTCTASGFSL----SNYYVTWVRQAPGKGLEWIGIIYGS---DETAYATWAI-GRFTISKTST--TVDLKMTSLTAADTATYFCARDDSSD-----------------------WDAKFNLWGQGTLVTVSS |
| Experimental | EVQLVESGG-GLVQPGGSLRLSCAASGFSL----SNYYVTWVRQAPGKGLEWVGIIYGS---DETAYATSAI-GRFTISRDNSKNTLYLQMNSLRAEDTAVYYCARDDSSD-----------------------WDAKFNLWGQGTLVTVSS |
| Hu-mAb | Q-SLEESGG-RLVTPGGSLRLTCAASGFSL----SNYYVTWVRQAPGKGLEWISIIYGS---DETAYATWAI-GRFTISKTST--TVDLKMTSLRAADTAVYFCARDDSSD-----------------------WDAKFNLWGQGTLVTVSS |

Light Chain

### experimental mutations: 22
### humab mutations: 4
mutation ratio: 18%
overlap ratio: 75%
adjusted OR: 75%

| | |
|---|---|
| Precursor | AYDMTQTPASVSAAVGGTVTIKCQASQSI------NNELSWYQQKPGQRPKLLIYRA-------STLASGVS-SRFKGSG--SGTEFTLTISDLECADAATYYCQQGYSL-------------------------RNIDNAFGGGTEVVVK- |
| Experimental | AIQMTQSPSSLSASVGDRVTITCQASQSI------NNELSWYQQKPGKAPKLLIYRA-------STLASGVP-SRFSGSG--SGTDFTLTISSLQPDDFATYYCQQGYSL-------------------------RNIDNAFGGGTKVEIK- |
| Hu-mAb | AYDMTQTPASVSAAVGGRVTIKCQASQSI------NNELSWYQQKPGQAPKLLIYRA-------STLQSGVS-SRFKGSG--SGTEFTLTISDLQCADAATYYCQQGYSL-------------------------RNIDNAFGGGTEVVVK- |

**Ligelizumab** Heavy Chain

### experimental mutations: 21
### humab mutations: 11
mutation ratio: 52%
overlap ratio: 64%
adjusted OR: 64%

| | |
|---|---|
| Precursor | QVQLQQSGA-ELMKPGASVKISCKTTGYTF----SMYWLEWVKQRPGHGLEWVGEISPG--TFTTNYNEKFK-AKATFTADTSSNTAYLQLSGLTSEDSAVYFCARFSHFSG---------------------SNYDYFDYWGQGTSLTVSS |
| Experimental | QVQLVQSGA-EVMKPGSSVKVSCKASGYTF----SWYWLEWVRQAPGHGLEWMGEIDPG--TFTTNYNEKFK-ARVTFTADTSTSTAYMELSSLRSEDTAVYYCARFSHFSG---------------------SNYDYFDYWGQGTLVTVSS |
| Hu-mAb | QVQLVQSGA-EVKKPGASVKVSCKTTGYTF----SMYWLEWVRQAPGHGLEWVGEISPG--TFTTNYAEKFQ-AKVTFTADTSINTAYLELSGLTSEDSAVYFCARFSHFSG---------------------SNYDYFDYWGQGTSLTVSS |

Light Chain

### experimental mutations: 21
### humab mutations: 11
mutation ratio: 52%
overlap ratio: 64%
adjusted OR: 91%

| | |
|---|---|
| Precursor | DILLTQSPAILSVSPGERVSFSCRASQSI------GTNIHWYQQRTDGSPRLLIKYA-------SESISGIP-SRFSGSG--SGTEFTLNINSVESEDIADYYCQQSDS----------------------------WPTTFGGGTKLEIK- |
| Experimental | EIVMTQSPATLSVSPGERATLSCRASQSI------GTNIHWYQQKPGQAPRLLIYYA-------SESISGIP-ARFSGSG--SGTEFTLTISSLQSEDFAVYYCQQSWS----------------------------WPTTFGGGTKVEIK- |
| Hu-mAb | EILLTQSPAILSVSPGERATFSCRASQSI------GTNLAWYQQRTDQSPRLLIYYA-------SESATGIP-ARFSGSG--SGTEFTLNINSVESEDIAVYYCQQSDS----------------------------WPTTFGGGTKLEIK- |

**Crizanlizumab** Heavy Chain

### experimental mutations: 23
### humab mutations: 11
mutation ratio: 48%
overlap ratio: 64%
adjusted OR: 64%

| | |
|---|---|
| Precursor | QVQLQQSGP-ELVKPGALVKISCKASGYTF----TSYDINWVKQRPGQGLEWIGWIYPG--DGSIKYNEKFK-GKATLTVDKSSSTAYMQVSSLTSENSAVYFCARRGEYGN----------------------YEGAMDYWGQGTTVTVSS |
| Experimental | QVQLVQSGA-EVKKPGASVKVSCKVSGYTF----TSYDINWVRQAPGKGLEWMGWIYPG--DGSIKYNEKFK-GRVTMTVDKSTDTAYMELSSLRSEDTAVYYCARRGEYGN----------------------YEGAMDYWGQGTLVTVSS |
| Hu-mAb | QVQLVQSGP-EVKKPGALVKISCKASGYTF----TSYDINWVRQAPGQGLEWMGWIYPG--DGSIKYAEKFQ-GKATLTVDKSISTAYMEVSSLTSENMAVYFCARRGEYGN----------------------YEGAMDYWGQGTTVTVSS |

Light Chain

### experimental mutations: 23
### humab mutations: 20
mutation ratio: 87%
overlap ratio: 85%
adjusted OR: 95%

| | |
|---|---|
| Precursor | DIVLTQSPASLAVSLGQRATISCKASQSVDY--DGHSYMNWYQQKPGQPPKLLIYAA-------SNLESGIP-ARFSGSG--SGTDFTLNIHPVEEEDAATYYCQQSDE----------------------------NPLTFGTGTKLELK- |
| Experimental | DIQMTQSPSSLSASVGDRVTITCKASQSVDY--DGHSYMNWYQQKPGKAPKLLIYAA-------SNLESGVP-SRFSGSG--SGTDFTLTISSLQPEDFATYYCQQSDE----------------------------NPLTFGGGTKVEIK- |
| Hu-mAb | DIQLTQSPSSLSASVGDRATITCQASQSVDY--DGHSYLNWYQQKPGKAPKLLIYAA-------SNLESGVP-SRFSGSG--SGTDFTLNIHSLQPEDFATYYCQQSDE----------------------------NPLTFGTGTRVELK- |

**Mogamulizumab** Heavy Chain

### experimental mutations: 15
### humab mutations: 6
mutation ratio: 40%
overlap ratio: 67%
adjusted OR: 67%

| | |
|---|---|
| Precursor | EVQLVESGG-DLMKPGGSLKISCAASGFIF----SNYGMSWVRQTPDMRLEWVATISSA--STYSYYPDSVK-GRFTISRDNAENSLYLQMNSLRSEDTGIYYCGRHSDG-------------------------NFAFGYWGRGTLVTVSA |
| Experimental | EVQLVESGG-DLVQPGRSLRLSCAASGFIF----SNYGMSWVRQAPGKGLEWVATISSA--STYSYYPDSVK-GRFTISRDNAKNSLYLQMNSLRVEDTALYYCGRHSDG-------------------------NFAFGYWGQGTLVTVSS |
| Hu-mAb | QVQLVESGG-DLMKPGGSLRISCAASGFIF----SNYGMSWVRQTPGMGLEWVATISSA--STYSYYADSVK-GRFTISRDNAENSLYLQMNSLRVEDTGIYYCGRHSDG-------------------------NFAFGYWGRGTLVTVSA |

Light Chain

### experimental mutations: 12
### humab mutations: 6
mutation ratio: 50%
overlap ratio: 67%
adjusted OR: 67%

| | |
|---|---|
| Precursor | DVLMTQTPLSLPVSLGDQASISCRSSRNIVHI-NGDTYLEWYLQRPGQSPKLLIYKV-------SNRFSGVP-DRFSGSG--SGTDFTLKISRVEAEDLGVYYCFQGSL----------------------------LPWTFGGGTRLEIR- |
| Experimental | DVLMTQSPLSLPVTPGEPASISCRSSRNIVHI-NGDTYLEWYLQKPGQSPQLLIYKV-------SNRFSGVP-DRFSGSG--SGTDFTLKISRVEAEDVGVYYCFQGSL----------------------------LPWTFGQGTKVEIK- |
| Hu-mAb | DVLMTQTPLSLPVTLGQPASISCRSSRNIVHI-NGDTYLEWYLQRPGQSPRLLIYKV-------SNRFSGVP-DRFSGSG--SGTDFTLKISRVEAEDVGVYYCFQGSL----------------------------LPWTFGQGTRLEIR- |

**Refanezumab** Heavy Chain

### experimental mutations: 17
### humab mutations: 15
mutation ratio: 88%
overlap ratio: 87%
adjusted OR: 87%

| | |
|---|---|
| Precursor | EIQLVQSGP-ELKKPGETNKISCKASGYTF----TNYGMNWVKQAPGKGLKWMGWINTY--TGEPTYADDFT-GRFAFSLETSASTAYLQISNLKNEDTATYFCARNPINYYGI------------------NYEGYVMDYWGQGTLVTVSS |
| Experimental | QVQLVQSGS-ELKKPGASVKVSCKASGYTF----TNYGMNWVRQAPGQGLEWMGWINTY--TGEPTYADDFT-GRFVFSLDTSVSTAYLQISSLKAEDTAVYYCARNPINYYGI------------------NYEGYVMDYWGQGTLVTVSS |
| Hu-mAb | QVQLVQSGS-ELKKPGASNKVSCKASGYTF----TNYGMNWVRQAPGKGLEWMGWINTY--TGEPTYAQGFT-GRFVFSLDTSASTAYLQISNLKAEDTAVYYCARNPINYYGI------------------NYEGYVMDYWGQGTLVTVSS |

Light Chain

### experimental mutations: 17
### humab mutations: 12
mutation ratio: 71%
overlap ratio: 92%
adjusted OR: 100%

| | |
|---|---|
| Precursor | NIMMTQSPSSLAVSAGEKVTMSCKSSHSVLYSSNQKNYLAWYQQKPGQSPKLLIYWA-------STRESGVP-DRFTGSG--SGTDFTLTIINVHTEDLAVYYCHQYL-----------------------------SSLTFGTGTKLEIK- |
| Experimental | DIVMTQSPDSLAVSLGERATINCKSSHSVLYSSNQKNYLAWYQQKPGQPPKLLIYWA-------STRESGVP-DRFSGSG--SGTDFTLTISSLQAEDVAVYYCHQYL-----------------------------SSLTFGQGTKLEIK- |
| Hu-mAb | DIVMSQSPDSLAVSLGERATINCKSSHSVLYSSNQKNYLAWYQQKPGQPPKLLIYWA-------STRESGVP-DRFTGSG--SGTDFTLTIINLHTEDVAVYYCHQYL-----------------------------SSLTFGTGTKLEIK- |
