## Supplementary information (main) for "Humanization of antibodies using a machine learning approach on large-scale repertoire data"

Contents:

1. Data
    A. Overview of sequences downloaded from the Observed Antibody Space database
    B. Therapeutics – sequences
    C. Therapeutics – origin species
    D. Therapeutics – ADA data
    E. Hu-mAb test set - references and reported immunogenicity

2. Methods
    A. 'Infixes' used for therapeutic classification
    B. Hu-mAb pipeline
    C. Residue type definitions for calculation of the Adjusted Overlap Ratio

3. Results
    A. PCA analysis of VH sequences by V gene and J gene type
    B. Classification Performance of RF models
    C. Comparison of our RF models with LSTM
    D. Classification of therapeutics using LSTM
    E. Relationship of RF humanness scores with immunogenicity
    F. Hu-mAb humanization results
    G. Random humanization results
    H. Analysis of proposed mutations – residue types
    I. Analysis of proposed mutations – residue locations

### 1. Data

#### A. Overview of sequences downloaded from the Observed Antibody Space database

Human sequences:

| | VH | VL (kappa) | VL (lambda) |
|---|---|---|---|
| V1 | 1,189,145 | 8,445,547 | 7,343,760 |
| V2 | 52,673 | 2,873,426 | 9,005,751 |
| V3 | 2,680,192 | 8,678,865 | 4,788,775 |
| V4 | 1,075,999 | 3,245,968 | 747,946 |
| V5 | 87,227 | 32,593 | 256,729 |
| V6 | 29,894 | 131,586 | 429,196 |
| V7 | 17,989 | | 637,720 |
| V8 | | | 409,564 |
| V10 | | | 32,503 |
| **Total** | **5,133,119** | **23,407,985** | **23,209,877** |

Negative (non-human) sequences:

| | VH | VL (kappa) | VL (lambda) |
|---|---|---|---|
| **Total** | **12,284,297** | **950,321** | **655,818** |

#### B. Therapeutics - sequences

For the list of 481 therapeutics and their sequences, please see the SI file 'Therapeutic_Sequences.xlsx'. The precursor and experimentally humanized sequences of the 25 therapeutics used to test Hu-mAb can be found in the SI file 'Hu-mAb_Results.pdf'.

#### C. Therapeutics - origin species

Therapeutic antibodies split by origin that are approved or in phase 1-3 trials. The therapeutics were gathered from the Therapeutic Structural Antibody Database, Thera-SAbDab (Raybould et al., 2020), which had a total of 481 antibody therapeutics intended for human use.

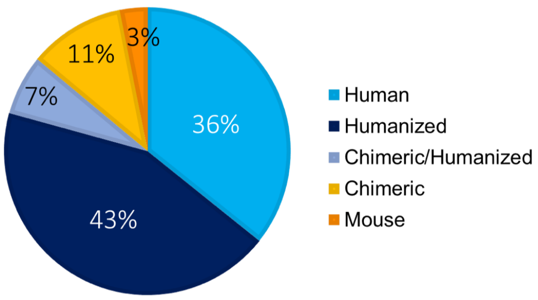


#### D. Therapeutics - ADA data

For the list of 481 therapeutics and their sequences, please see the SI file 'Therapeutic_ADA.xlsx'.

### E. Hu-mAB test set

25 therapeutics were selected for which both the precursor and experimentally humanized sequences were available. The references for these sequences are as follows:

| Therapeutic | Reference |
|---|---|
| AntiCD28 | https://www.jimmunol.org/content/169/2/1119 |
| Campath | https://journals.lww.com/transplantjournal/Fulltext/1999/11150/ANTI_GLOBULIN_RESPONSES_TO_RAT_AND_HUMANIZED.32.aspx |
| Bevacizumab | http://www.imgt.org/IMGTrepertoire/GenesClinical/humanized/bevacizumab/bevacizumab_ProteinDisplay.html#igh |
| Herceptin | https://www.ncbi.nlm.nih.gov/pmc/articles/PMC49066/pdf/pnas01084-0075.pdf |
| Omalizumab | https://www.jimmunol.org/content/151/5/2623 |
| Eculizumab | https://patentimages.storage.googleapis.com/f2/c4/09/171125042450cd/EP2298808A1.pdf |
| Tocilizumab | https://cancerres.aacrjournals.org/content/canres/53/4/851.full.pdf |
| Pembrolizumab | https://cancerres.aacrjournals.org/content/74/19_Supplement/5024 |
| Pertuzumab | https://pubmed.ncbi.nlm.nih.gov/16151804/ |
| Ixekizumab | https://www.ncbi.nlm.nih.gov/pmc/articles/PMC4846058/#SD1-jir-9-039 |
| Palivizumab | https://academic.oup.com/jid/article/176/5/1215/831423 |
| Certolizumab | https://patentimages.storage.googleapis.com/54/71/03/400fe464c8bb2d/US20050042219A1.pdf |
| Idarucizumab | https://patentimages.storage.googleapis.com/51/ff/74/48026e9919861a/EP2525812B1.pdf |
| Reslizumab | https://patentimages.storage.googleapis.com/b1/be/fc/ee66a606c6ed4f/CA2192543C.pdf |
| Solanezumab | https://patentimages.storage.googleapis.com/8a/ed/d6/645d49a2b2fae4/WO2004071408A2.pdf |
| Lorvotuzumab | https://patentimages.storage.googleapis.com/6e/13/b2/f740eceb58298a/US5639641.pdf |
| Pinatuzumab | https://patentimages.storage.googleapis.com/e1/b4/6f/b94b77b6b5806f/ES2543475T3.pdf |
| Etaracizumab | https://www.pnas.org/content/pnas/95/15/8910.full.pdf |
| Talacotuzumab | https://patentimages.storage.googleapis.com/74/d1/05/9d3a61813b2985/US8492119.pdf |
| Rovalpituzumab | https://patentimages.storage.googleapis.com/4a/00/25/e01f76b1cb6ec6/US9089616.pdf |
| Clazakizumab | https://patentimages.storage.googleapis.com/52/b8/4f/0146181ade3705/US20090104187A1.pdf |
| Ligelizumab | https://patentimages.storage.googleapis.com/9d/5c/b5/f8789f9a5c7722/US7531169.pdf |
| Crizanlizumab | https://patentimages.storage.googleapis.com/b9/22/2d/5e02e51d7e935a/US8377440.pdf |
| Mogamulizumab | https://patentimages.storage.googleapis.com/20/fc/8e/b206ab42434698/US8491902.pdf |
| Refanezumab | https://patentimages.storage.googleapis.com/1b/f7/0c/0e94c7d7cf18ad/US8974782.pdf |

Reported immunogenicity for these 25 therapeutics is shown in the table below. All therapeutics except for two (Talacotuzumab/AntiCD28) have low immunogenicity (<10% patients with ADA). Immunogenicity of Anti-CD28 was measured *in vitro* using a mixed lymphocyte reaction (MLR) assay and the precursor sequence was found to have a much higher immunogenic response compared to the experimental humanized sequence.

| Therapeutic | Immunogenicity (% of patients with ADA) |
|---|---|
| AntiCD28 | NA |
| Campath | 5.10 |
| Bevacizumab | 0.32 |
| Herceptin | 8.10 |
| Omalizumab | 0.00 |
| Eculizumab | 2.00 |
| Tocilizumab | 2.00 |
| Pembrolizumab | 1.70 |
| Pertuzumab | 2.80 |
| Ixekizumab | 8.50 |
| Palivizumab | 1.10 |
| Certolizumab | 8.00 |
| Idarucizumab | 4.00 |
| Reslizumab | 5.00 |
| Solanezumab | 3.50 |
| Lorvotuzumab | 0.00 |
| Pinatuzumab | 1.40 |
| Etaracizumab | 0.00 |
| Talacotuzumab | 17.40 |
| Rovalpituzumab | 0.00 |
| Clazakizumab | 1.82 |
| Ligelizumab | 6.32 |
| Crizanlizumab | 0.94 |
| Mogamulizumab | 4.20 |
| Refanezumab | 9.38 |

### 2. Methods

#### A. 'Infixes' used for therapeutic classification

| Source infix | Origin |
|---|---|
| -u- | Human |
| -zu- | Humanized |
| -xizu- | Chimeric/Humanized |
| -xi- | Chimeric |
| -o- | Mouse |

#### B. Hu-mAb pipeline

Outline of the Hu-mAb protocol. All possible mutations within the framework region of the input sequence were made and each sequence scored by the respective RF model. The mutated sequence with the highest humanness score was selected and the process repeated until the target score was achieved.

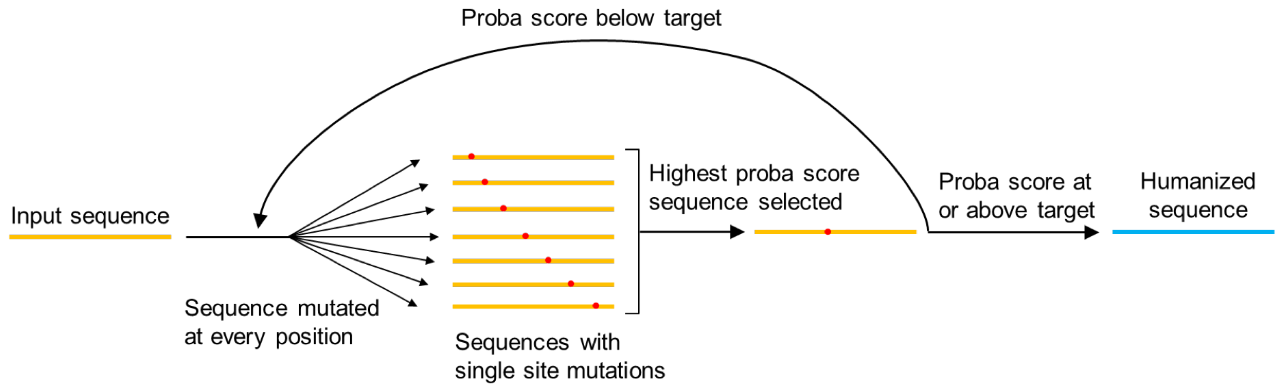


#### C. Residue type definitions for calculation of the Adjusted Overlap Ratio

Groupings of amino acid types used to calculate the Adjusted Overlap Ratio. Adjusted Overlap Ratio (AOR) treats any mutation to the same type as an overlapping / identical mutation. For example, if the experimental mutation was R to A and the model mutation was R to V, this would still be considered an overlap. Others are not included as part of the AOR calculation and each AA is treated separately as a separate type.

| Type | Amino Acids |
|---|---|
| Positive | K, R, H |
| Negative | D, E |
| Hydrophobic | V, M, I, L, A |
| Hydrophilic | Q, N, S, T |
| Aromatic | W, F, Y |
| Others | C, G, P |

### 3. Results

#### A. PCA analysis of VH sequences by V gene and J gene type

Prior to model construction the dataset was split to improve classification performance and prevent unrealistic humanization suggestions. The variable domain of each antibody is predominantly made from a single V gene and it was expected that sequences from the same V gene would be more similar than sequences from other V genes. Moreover, the humanizer was envisaged to propose mutations towards a particular V gene type rather than an unrealistic hybrid sequence. PCA was performed to test for the dissimilarity across sequences from different V gene types. All human IgG VH sequences were downloaded from OAS (Kovaltsuk et al., 2018) and prepared as described in Methods. PCA was performed and the sequences were labelled by their V gene types (V1-V7) (SI Figure 1). Each set of V gene sequences demonstrated trivial clustering and were completely separable from all other sets of V gene sequences with just two or three principal components. The same approach above was performed with J genes. Each set of V gene sequences were split into their respective J gene types (J1-J6). PCA was performed on each set of V gene sequences and were labelled by their J gene type. Figure 1E shows no distinct clustering across the V3 J gene sequences and were completely inseparable across all three principal components. The lack of separability by J gene type was consistent across all sets of V gene sequences. Therefore, sequences were only split by V gene and not J gene type for model construction.

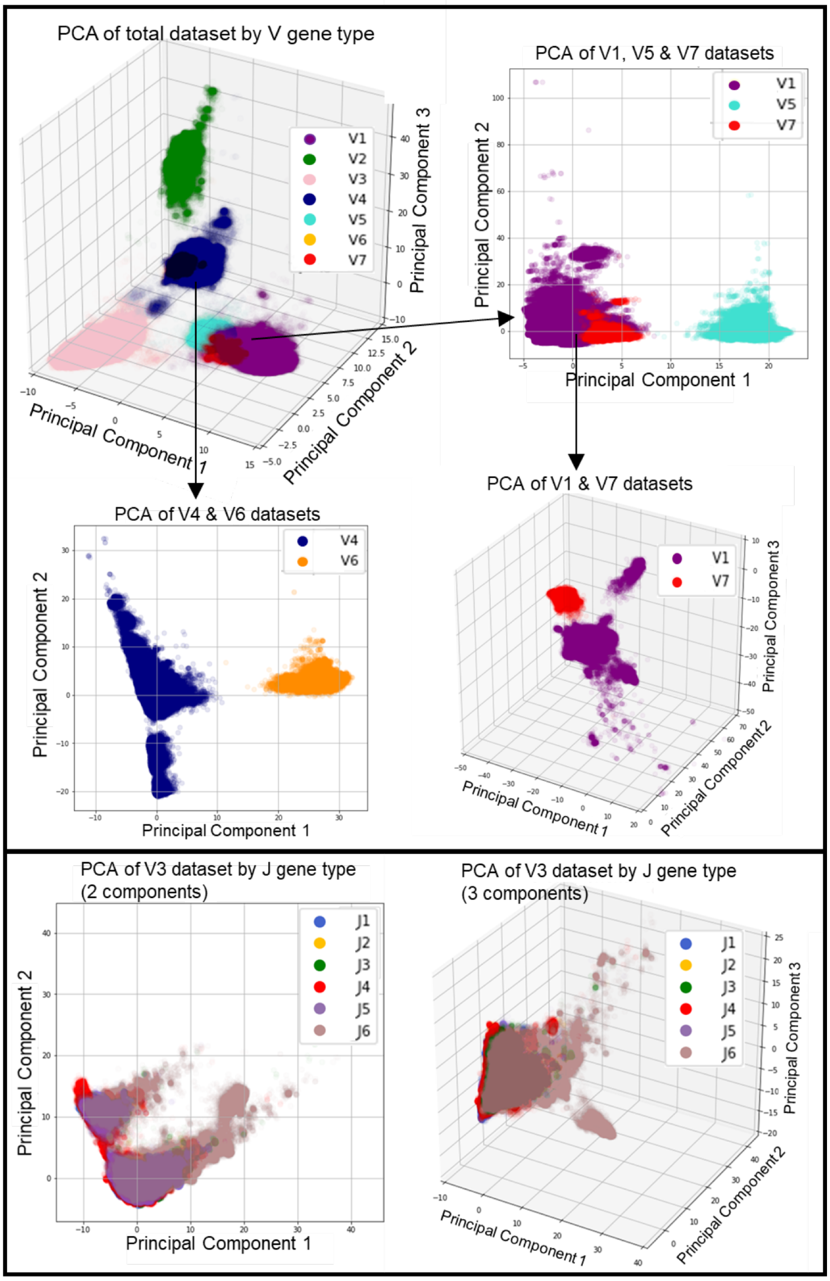


Principal Component Analysis of VH sequences by V gene type and J gene type. Sequences were labelled by their V gene type (A,B,C,D) or by their J gene type (E). A) Three component PCA of all VH sequences exhibited four distinct clusters of sequences; consisting of (i) V2, (ii) V3, (iii) V4/V6 and (iv) V1/V5/V7. Further PCA was performed independently on sequences in cluster (iii) and (iv). B) Two component PCA of V4 and V6 sequences exhibited distinct clustering of each set. C) Two component PCA of V1, V5 and V7 sequences resulted in two distinct clusters; V1/V7 and V5. Further PCA was performed on just V1 and V7. D) Three component PCA of V1 and V7 sequences exhibited distinct clustering of each set. E) Two component and three component PCA of V3 sequences labelled by their J gene type. No distinct clustering was observed with all sets of J gene sequences overlapped.

### B. Classification performance of RF models

Validation and Test Set performance for each RF model. Extremely high performance was observed across all models demonstrating AUC scores close to 1. The VH models were able to perfectly discriminate between human and mouse sequences in some cases with AUC scores of 1. YJS scores were similar in both validation and test sets with all models scoring ≥0.999. MCC thresholds obtained using the validation set were very similar to the YJS thresholds - the YJS thresholds were taken forward for use in subsequent experiments.

Validation:

| | V Gene | ROCAUC | YJS | YJS Threshold | MCC | MCC Threshold |
|---|---|---|---|---|---|---|
| HEAVY | HV1 | 1.0000000000 | 1.000000 | 0.565 | 1.000000 | 0.565 |
| | HV2 | 1.0000000000 | 1.000000 | 0.615 | 1.000000 | 0.615 |
| | HV3 | 1.0000000000 | 1.000000 | 0.630 | 1.000000 | 0.630 |
| | HV4 | 1.0000000000 | 1.000000 | 0.495 | 1.000000 | 0.495 |
| | HV5 | 1.0000000000 | 1.000000 | 0.480 | 1.000000 | 0.480 |
| | HV6 | 1.0000000000 | 1.000000 | 0.640 | 1.000000 | 0.640 |
| | HV7 | 1.0000000000 | 1.000000 | 0.575 | 1.000000 | 0.575 |
| KAPPA | KV1 | 0.9999990912 | 0.999906 | 0.740 | 0.999538 | 0.714 |
| | KV2 | 0.9999999990 | 0.999986 | 0.602 | 0.999972 | 0.642 |
| | KV3 | 0.9999999989 | 0.999977 | 0.845 | 0.999737 | 0.722 |
| | KV4 | 0.9999999998 | 0.999994 | 0.650 | 0.999993 | 0.650 |
| | KV5 | 1.0000000000 | 1.000000 | 0.515 | 1.000000 | 0.515 |
| | KV6 | 1.0000000000 | 1.000000 | 0.490 | 1.000000 | 0.490 |
| LAMBDA | LV1 | 0.9999999999 | 0.999992 | 0.856 | 0.999988 | 0.816 |
| | LV2 | 0.9999999996 | 0.999997 | 0.790 | 0.999984 | 0.772 |
| | LV3 | 0.9999999990 | 0.999967 | 0.770 | 0.999965 | 0.800 |
| | LV4 | 0.9999999990 | 0.999973 | 0.815 | 0.999986 | 0.800 |
| | LV5 | 0.9999999994 | 0.999985 | 0.370 | 0.999973 | 0.510 |
| | LV6 | 1.0000000000 | 1.000000 | 0.800 | 1.000000 | 0.800 |
| | LV7 | 1.0000000000 | 1.000000 | 0.810 | 1.000000 | 0.810 |
| | LV8 | 0.9999999985 | 0.999985 | 0.754 | 0.999990 | 0.754 |
| | LV10 | 1.0000000000 | 1.000000 | 0.625 | 1.000000 | 0.590 |

Testing:

| | V Gene | ROCAUC | YJS | MCC |
|---|---|---|---|---|
| HEAVY | HV1 | 1.00000000000 | 1.000000 | 1.000000 |
| | HV2 | 1.00000000000 | 1.000000 | 1.000000 |
| | HV3 | 1.00000000000 | 1.000000 | 1.000000 |
| | HV4 | 1.00000000000 | 1.000000 | 1.000000 |
| | HV5 | 1.00000000000 | 1.000000 | 1.000000 |
| | HV6 | 1.00000000000 | 1.000000 | 1.000000 |
| | HV7 | 1.00000000000 | 1.000000 | 1.000000 |
| KAPPA | KV1 | 0.99999981855 | 0.999552 | 0.999540 |
| | KV2 | 0.99999999844 | 0.999958 | 0.999970 |
| | KV3 | 0.99999999770 | 0.999526 | 0.999740 |
| | KV4 | 0.99999999998 | 0.999997 | 0.999990 |
| | KV5 | 1.00000000000 | 1.000000 | 1.000000 |
| | KV6 | 1.00000000000 | 1.000000 | 1.000000 |
| LAMBDA | LV1 | 0.99999999994 | 0.999996 | 0.999980 |
| | LV2 | 0.99999999998 | 0.999997 | 0.999980 |
| | LV3 | 0.99999998860 | 0.999950 | 0.999960 |
| | LV4 | 1.00000000000 | 0.999987 | 0.999990 |
| | LV5 | 0.99999999941 | 0.999995 | 0.999920 |
| | LV6 | 1.00000000000 | 1.000000 | 1.000000 |
| | LV7 | 1.00000000000 | 1.000000 | 1.000000 |
| | LV8 | 1.00000000000 | 1.000000 | 1.000000 |
| | LV10 | 1.00000000000 | 0.999692 | 0.999840 |

### C. Comparison of our RF models with LSTM

Our RF models outperform the LSTM equivalents in both AUC and YJS scores for each model.

| | | Random Forest | | LSTM | |
|---|---|---|---|---|---|
| | V Gene | ROCAUC | YJS | ROCAUC | YJS |
| HEAVY | HV1 | 1.0000000000 | 1.000000 | 0.999772 | 0.9960 |
| | HV2 | 1.0000000000 | 1.000000 | 0.999996 | 0.9970 |
| | HV3 | 1.0000000000 | 1.000000 | 0.994383 | 0.9418 |
| | HV4 | 1.0000000000 | 1.000000 | 0.991764 | 0.9917 |
| | HV5 | 1.0000000000 | 1.000000 | 0.999954 | 0.9981 |
| | HV6 | 1.0000000000 | 1.000000 | 0.999999 | 0.9997 |
| | HV7 | 1.0000000000 | 1.000000 | 0.999991 | 0.9991 |
| KAPPA | KV1 | 0.9999998186 | 0.999552 | 0.939153 | 0.6790 |
| | KV2 | 0.9999999984 | 0.999958 | 0.997548 | 0.9481 |
| | KV3 | 0.9999999977 | 0.999526 | 0.993947 | 0.9156 |
| | KV4 | 1.0000000000 | 0.999997 | 0.998431 | 0.9746 |
| | KV5 | 1.0000000000 | 1.000000 | 0.999992 | 0.9993 |
| | KV6 | 1.0000000000 | 1.000000 | 0.999683 | 0.9930 |
| LAMBDA | LV1 | 0.9999999999 | 0.999996 | 0.998347 | 0.9702 |
| | LV2 | 1.0000000000 | 0.999997 | 0.995076 | 0.9191 |
| | LV3 | 0.9999999886 | 0.999950 | 0.999284 | 0.9740 |
| | LV4 | 1.0000000000 | 0.999987 | 0.999989 | 0.9989 |
| | LV5 | 0.9999999994 | 0.999995 | 0.999981 | 0.9959 |
| | LV6 | 1.0000000000 | 1.000000 | 0.999962 | 0.9939 |
| | LV7 | 1.0000000000 | 1.000000 | 0.999802 | 0.9919 |
| | LV8 | 1.0000000000 | 1.000000 | 0.999999 | 0.9996 |
| | LV10 | 1.0000000000 | 0.999692 | 0.999732 | 0.9933 |

### D. Classification of Therapeutics using LSTM

Percentage of antibody therapeutics classified as human by the LSTM method, split by their origin. Chi/humanized are sequences which are part humanized andpart chimeric. Therapeutics were classified based on their VH and VL sequences separately, as well as combined (to be classified as human, both VH and VL scores had to be above the respective humanness threshold). For our RF models, the predicted humanness of the therapeutics decreased as the human content of the sequence decreases (left to right in the figure). This trend is also observed using the LSTM method, but not as clearly - for example, more of the chi/humanized set are classified as human than the humanized set, and some human therapeutics are classified as non-human.

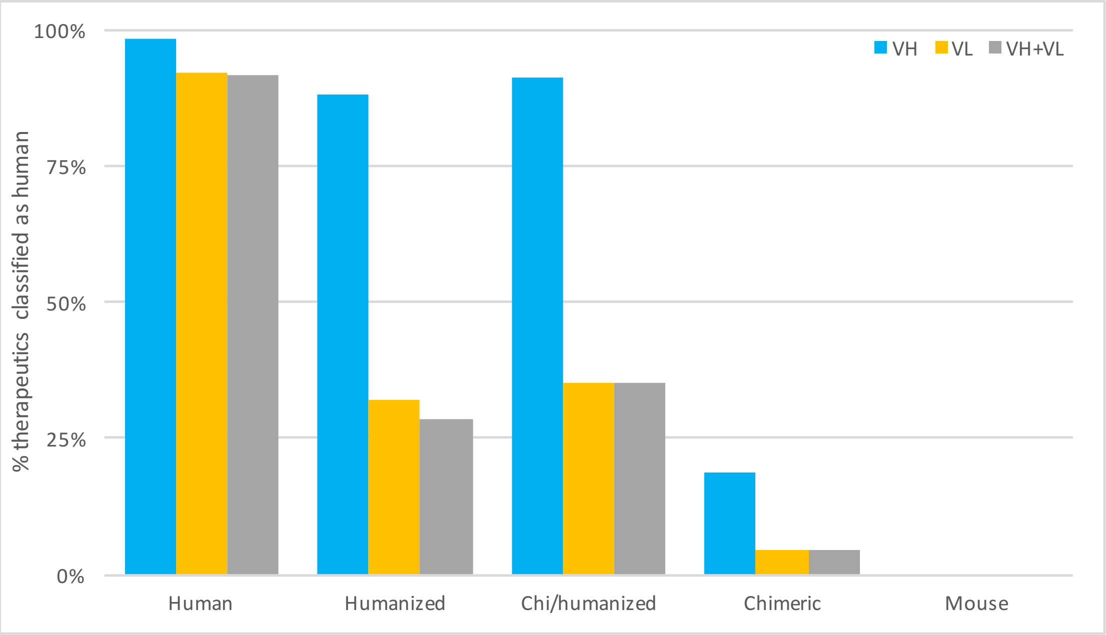

#### E. Relationship of RF humanness scores with immunogenicity

Scatter plot comparing experimental immunogenicity (fraction of patients that develop ADAs) and the RF humanness score of therapeutic mAbs. A) VL sequence. B) VH sequences. C) Combined sequences – the average proba score of the respective VH and VL sequence.

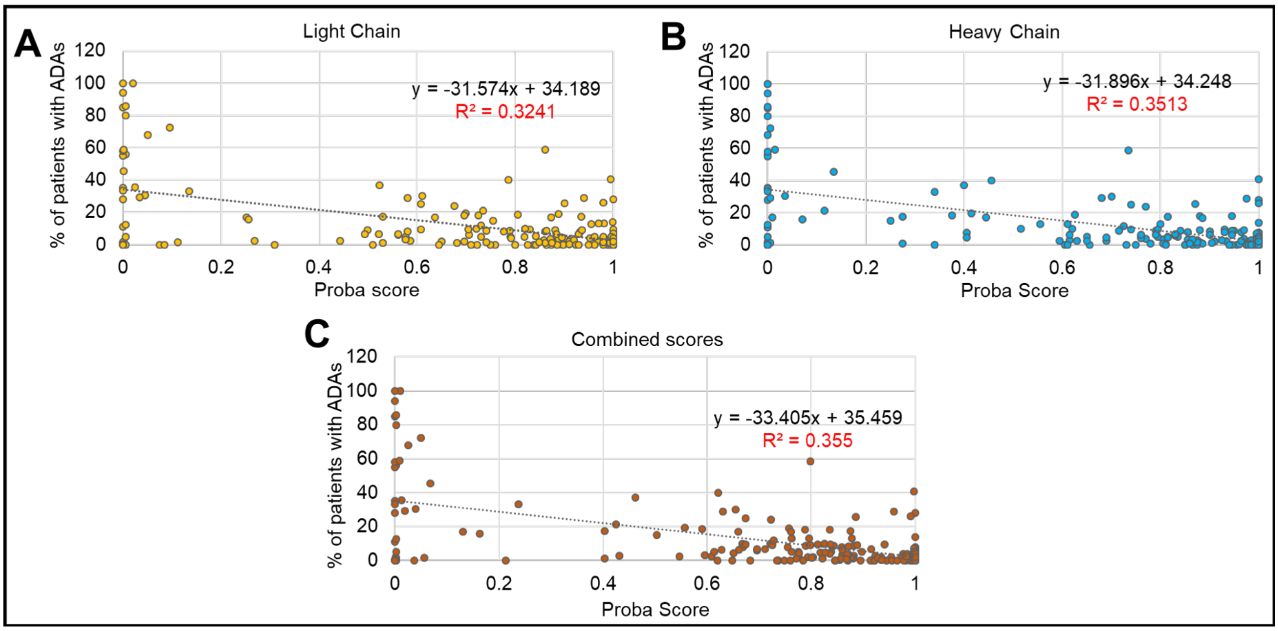


#### F. Hu-mAb humanization results

For full results, please see SI file 'Hu-mAb_Results.pdf'. This file contains the precursor, experimentally humanized and Hu-mAb output sequences for each of the 25 therapeutics.

Heavy chains summary:

| Therapeutic | V Gene | Initial Score | Target Score | # Exp. Mutations | # Hu-mAb Mutations | Mutation Ratio | Overlap Ratio | Adjusted OR |
|---|---|---|---|---|---|---|---|---|
| AntiCD28 | hv3 | 0.005 | 0.990 | 33 | 19 | 58% | 63% | 79% |
| Campath | hv4 | 0.000 | 0.835 | 39 | 16 | 41% | 75% | 88% |
| Bevacizumab | hv3 | 0.050 | 0.805 | 25 | 14 | 56% | 50% | 57% |
| Herceptin | hv3 | 0.000 | 0.965 | 32 | 27 | 84% | 59% | 78% |
| Omalizumab | hv3 | 0.000 | 0.955 | 34 | 21 | 62% | 62% | 76% |
| Eculizumab | hv1 | 0.000 | 0.975 | 23 | 15 | 65% | 73% | 73% |
| Tocilizumab | hv4 | 0.000 | 0.935 | 23 | 14 | 61% | 64% | 86% |
| Pembrolizumab | hv1 | 0.000 | 0.880 | 23 | 11 | 48% | 73% | 73% |
| Pertuzumab | hv3 | 0.000 | 0.875 | 32 | 19 | 59% | 68% | 79% |
| Ixekizumab | hv1 | 0.000 | 0.900 | 29 | 12 | 41% | 75% | 75% |
| Palivizumab | hv2 | 0.000 | 0.885 | 18 | 12 | 67% | 75% | 83% |
| Certolizumab | hv3 | 0.000 | 0.855 | 31 | 18 | 58% | 61% | 78% |
| Idarucizumab | hv4 | 0.000 | 0.760 | 24 | 15 | 63% | 80% | 80% |
| Reslizumab | hv3 | 0.060 | 0.760 | 21 | 10 | 48% | 50% | 80% |
| Solanezumab | hv3 | 0.005 | 0.855 | 16 | 10 | 63% | 50% | 70% |
| Lorvotuzumab | hv3 | 0.010 | 0.985 | 13 | 10 | 77% | 90% | 90% |
| Pinatuzumab | hv3 | 0.000 | 0.860 | 33 | 23 | 70% | 61% | 78% |
| Etaracizumab | hv3 | 0.005 | 0.940 | 16 | 12 | 75% | 58% | 83% |
| Talacotuzumab | hv5 | 0.000 | 0.815 | 33 | 18 | 55% | 78% | 83% |
| Rovalpituzumab | hv1 | 0.000 | 0.975 | 30 | 21 | 70% | 67% | 67% |
| Clazakizumab | hv3 | 0.645 | 0.995 | 27 | 7 | 26% | 86% | 86% |
| Ligelizumab | hv1 | 0.000 | 0.805 | 21 | 11 | 52% | 64% | 64% |
| Crizanlizumab | hv1 | 0.000 | 0.865 | 23 | 11 | 48% | 64% | 64% |
| Mogamulizumab | hv3 | 0.050 | 0.760 | 15 | 6 | 40% | 67% | 67% |
| Refanezumab | hv7 | 0.025 | 0.860 | 17 | 15 | 88% | 87% | 87% |

| Therapeutic | V Gene | Initial Score | Target Score | # Exp. Mutations | # Hu-mAb Mutations | Mutation Ratio | Overlap Ratio | Adjusted OR |
|---|---|---|---|---|---|---|---|---|
| **MEAN** | | | | | | **59%** | **68%** | **77%** |
| **MEDIAN** | | | | | | **59%** | **67%** | **78%** |

Light chains summary:

| Therapeutic | V Gene | Initial Score | Target Score | # Exp. Mutations | # Hu-mAb Mutations | Mutation Ratio | Overlap Ratio | Adjusted OR |
|---|---|---|---|---|---|---|---|---|
| AntiCD28 | kv4 | 0.050 | 0.805 | 19 | 11 | 58% | 64% | 73% |
| Campath | kv1 | 0.724 | 0.842 | 14 | 3 | 21% | 67% | 67% |
| Bevacizumab | kv1 | 0.017 | 0.899 | 16 | 9 | 56% | 89% | 100% |
| Herceptin | kv1 | 0.032 | 0.776 | 22 | 8 | 36% | 88% | 88% |
| Omalizumab | kv1 | 0.081 | 0.874 | 25 | 19 | 76% | 89% | 95% |
| Eculizumab | kv1 | 0.002 | 0.893 | 20 | 12 | 60% | 83% | 83% |
| Tocilizumab | kv1 | 0.001 | 0.650 | 19 | 9 | 47% | 78% | 89% |
| Pembrolizumab | kv3 | 0.010 | 0.870 | 20 | 12 | 60% | 75% | 75% |
| Pertuzumab | kv1 | 0.006 | 0.888 | 20 | 10 | 50% | 80% | 90% |
| Ixekizumab | kv2 | 0.000 | 0.864 | 12 | 9 | 75% | 78% | 100% |
| Palivizumab | kv1 | 0.199 | 0.876 | 26 | 13 | 50% | 77% | 92% |
| Certolizumab | kv1 | 0.011 | 0.862 | 20 | 10 | 50% | 80% | 90% |
| Idarucizumab | kv2 | 0.258 | 0.900 | 8 | 6 | 75% | 67% | 67% |
| Reslizumab | kv1 | 0.389 | 0.791 | 20 | 6 | 30% | 83% | 100% |
| Solanezumab | kv2 | 0.060 | 0.888 | 10 | 8 | 80% | 88% | 100% |
| Lorvotuzumab | kv2 | 0.050 | 0.924 | 13 | 11 | 85% | 82% | 82% |
| Pinatuzumab | kv1 | 0.002 | 0.741 | 23 | 19 | 83% | 74% | 79% |
| Etaracizumab | kv3 | 0.010 | 0.950 | 25 | 13 | 52% | 62% | 69% |
| Talacotuzumab | kv4 | 0.005 | 0.935 | 16 | 11 | 69% | 73% | 73% |
| Rovalpituzumab | kv3 | 0.000 | 0.980 | 26 | 14 | 54% | 64% | 79% |
| Clazakizumab | kv1 | 0.778 | 0.911 | 22 | 4 | 18% | 75% | 75% |
| Ligelizumab | kv3 | 0.020 | 0.930 | 21 | 11 | 52% | 64% | 91% |
| Crizanlizumab | kv1 | 0.001 | 0.875 | 23 | 20 | 87% | 85% | 95% |
| Mogamulizumab | kv2 | 0.036 | 0.792 | 12 | 6 | 50% | 67% | 67% |
| Refanezumab | kv4 | 0.000 | 1.000 | 17 | 12 | 71% | 92% | 100% |
| **MEAN** | | | | | | **58%** | **77%** | **85%** |
| **MEDIAN** | | | | | | **56%** | **78%** | **88%** |

### G. Random humanization results

Random humanization of selected therapeutics. A random humanization model was constructed to generate mutations randomly up to the same number of mutations as Hu-mAb. 100 million randomly humanized VH sequences were generated and the average Overlap Ratios and Adjusted Overlap Ratios were calculated.

| Therapeutic | Overlap Ratio | Adjusted Overlap Ratio |
|---|---|---|
| Certolizumab | 1.8% | 6.0% |
| Omalizumab | 1.9% | 6.8% |
| Eculizumab | 1.3% | 5.2% |

#### H. Analysis of proposed mutations – residue types

Analysis of all mutations proposed by Hu-mAb and experiment using amino acid groupings given in section 2C. Also shown are the results of performing random mutations on the test sequences (the same number of mutations as proposed by Hu-mAb, repeated 1,000,000 times).

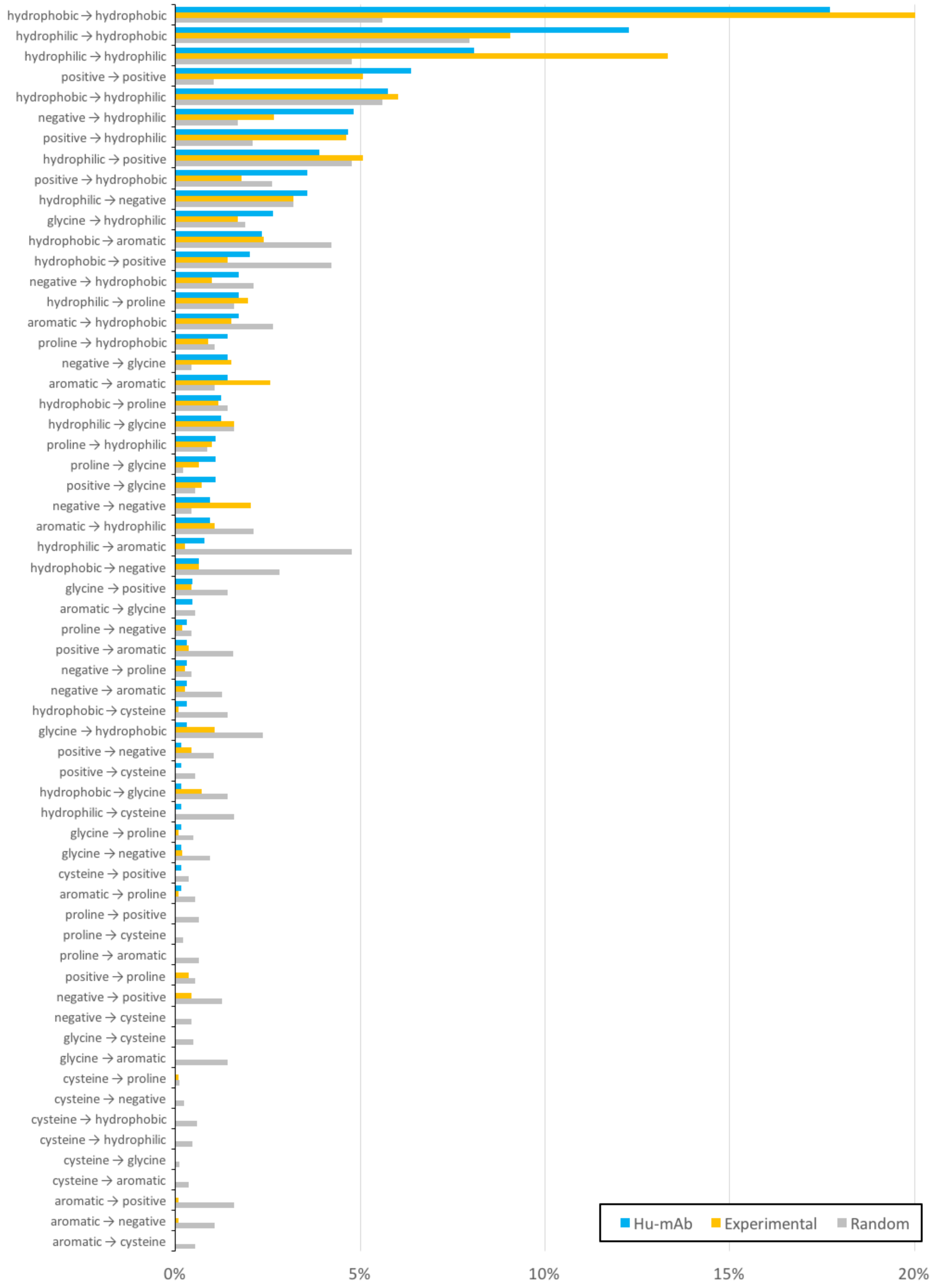

### I. Analysis of proposed mutations - residue locations

Comparison of mutation locations for humanizations performed experimentally and by Hu-mAb.

| | | VH | | | | | VL | | | | |
|---|---|---|---|---|---|---|---|---|---|---|---|
| | | Proportion of Mutations | | Mutations per Sequence | | Overlap Ratio | Proportion of Mutations | | Mutations per Sequence | | Overlap Ratio |
| Residues | | Hu-mAb | Experiment | Hu-mAb | Experiment | | Hu-mAb | Experiment | Hu-mAb | Experiment | |
| Interface | Mean | 6.2% | 7.3% | 0.8 | 1.6 | 73.8% | 8.2% | 10.1% | 0.8 | 1.8 | 96.4% |
| | Median | 4.3% | 5.7% | 1.0 | 2.0 | 100.0% | 7.7% | 10.0% | 1.0 | 2.0 | 100.0% |
| Vernier Zone | Mean | 14.1% | 10.4% | 2.2 | 3.0 | 51.7% | 4.8% | 5.0% | 0.4 | 1.0 | 70.0% |
| | Median | 14.3% | 12.5% | 2 | 3.0 | 58.3% | 0.0% | 4.6% | 0.0 | 1.0 | 100.0% |
| Surface | Mean | 67.1% | 67.6% | 9.6 | 17.1 | 72.5% | 63.3% | 64.9% | 6.7 | 12.4 | 78.1% |
| | Median | 66.7% | 66.7% | 9.0 | 16.0 | 70.0% | 63.6% | 66.7% | 6.0% | 13.0% | 83.3% |
| Buried | Mean | 32.8% | 29.8% | 5.1 | 8.1 | 51.5% | 36.7% | 32.6% | 3.9 | 6.3 | 74.3% |
| | Median | 33.3% | 33.3% | 5.0 | 9.0 | 60.0% | 36.4% | 31.3% | 4.0 | 7.0 | 75.0% |

Definitions of interface, Vernier zone, surface, and buried residues (IMGT residue numbering):

| Type | VH | VL |
|---|---|---|
| Interface | 44, 47, 48, 52, 101, 107, 109, 114, 116, 117, 120 | 50, 56, 69, 101, 103, 109, 115, 116, 120 |
| Vernier Zone | 2, 52, 53, 54, 76, 78, 80, 82, 87, 118 | 2, 4, 41, 42, 52, 53, 54, 55, 78, 80, 84, 85, 87, 118 |
| Surface | 1, 3, 5, 7, 8, 9, 11, 12, 14, 15, 16, 17, 18, 20, 22, 24, 26, 45, 46, 47, 48, 49, 51, 66, 69, 70, 72, 73, 74, 77, 79, 81, 82, 83, 84, 85, 88, 90, 92, 93, 95, 96, 97, 99, 101, 120, 123, 127, 128 | 1, 3, 5, 7, 8, 9, 10, 11, 12, 14, 15, 16, 17, 18, 20, 22, 24, 26, 45, 46, 47, 48, 51, 66, 67, 69, 70, 72, 73, 74, 77, 79, 80, 81, 82, 83, 84, 85, 86, 88, 90, 92, 93, 95, 96, 97, 101, 120, 123, 127, 128 |
| Buried | 2, 4, 6, 10, 13, 19, 21, 23, 25, 39, 40, 41, 42, 43, 44, 50, 52, 53, 54, 55, 67, 68, 71, 75, 76, 78, 80, 86, 87, 89, 91, 94, 98, 100, 102, 103, 104, 118, 119, 121, 122, 124, 125, 126 | 2, 4, 6, 13, 19, 21, 23, 25, 39, 40, 41, 42, 43, 44, 49, 50, 52, 53, 54, 55, 68, 71, 75, 76, 78, 87, 89, 91, 94, 98, 99, 100, 102, 103, 104, 118, 119, 121, 122, 124, 125, 126 |

Key interface residues were defined according to Raybould et al., 2020.

Vernier zone residues were defined according to Foote and Winter, 1992, converting the numbering scheme from Kabat to IMGT. Residue numbers 28-35 and 105-106 were excluded from the calculation since they are considered part of the CDRs according to the IMGT definition, and hence Hu-mAb would never propose mutations for those residues.

Surface/buried residues were defined using a set of 1129 non-redundant variable domain structures (from Raybould et al., 2020). We calculated the average relative solvent accessibility (RSA) for each position within the structures, and then set a threshold to split the residues between surface exposed and buried. This was defined as 25% of the RSA value, a value which is commonly used for other models (e.g. Wu et al., 2017, Zhang et al., 2017, Bozic et al., 2017).
